# Structural Basis of HIV-1 Maturation Inhibitor Binding and Activity

**DOI:** 10.1101/2022.02.22.481470

**Authors:** Sucharita Sarkar, Kaneil K. Zadrozny, Roman Zadorozhnyi, Ryan W. Russell, Caitlin M. Quinn, Alex Kleinpeter, Sherimay Ablan, Chaoyi Xu, Juan R. Perilla, Eric O. Freed, Barbie K. Ganser-Pornillos, Owen Pornillos, Angela M. Gronenborn, Tatyana Polenova

## Abstract

HIV-1 maturation inhibitors (MIs) interfere with the final step in the viral lifecycle by disrupting the ordered proteolytic processing of the viral Gag polyprotein into its individual domains. Bevirimat (BVM) and its analogs interfere with the final catalytic cleavage of spacer peptide 1 (SP1) from the capsid protein (CA) C-terminal domain (CA_CTD_), by binding to and stabilizing the CA_CTD_-SP1 region. MIs are under development as alternative drugs to augment current antiretroviral therapies. Although promising, their mechanism of action and associated virus resistance pathways remain poorly understood at the molecular, biochemical, and structural levels. Here, we report atomic-resolution magic angle spinning (MAS) NMR structures of microcrystalline assemblies of CA_CTD_-SP1 complexed with BVM and/or the assembly cofactor inositol hexakisphosphate (IP6). BVM and IP6 can bind simultaneously to SP1, with BVM positioned in the center of its 6-helix bundle in a unique conformation. Importantly, the NMR-observed structural effects of BVM on IP6 binding suggest that the inhibitor stabilizes the 6-helix bundle in multiple ways. In addition, BVM-resistant SP1-A1V and SP1-V7A variants exhibit distinct conformational and binding characteristics. Taken together, our results reveal a novel allosteric mechanism by which BVM disrupts maturation and provide a structural explanation for BVM resistance as well as important guidance for the design of new MIs.

## MAIN

HIV-1 maturation is triggered by the viral protease that cleaves the structural Gag polyprotein precursor into its constituent domains^1-3^. HIV-1 Gag harbors five proteolytic cleavage sites between its four major structural and functional domains and two spacer peptides: MA, CA, SP1, NC, SP2, and p6. Processing proceeds at different rates, with the SP1-NC site cleaved the fastest and the CA-SP1 site last^4^. Genetic and enzymatic studies showed that inhibition of cleavage or even slowing cleavage at the CA-SP1 site is sufficient to significantly disrupt the maturation process and abrogate virus infectivity. Indeed, maturation inhibitors (MIs) that interfere with CA-SP1 processing are emerging as attractive candidates for augmenting the current arsenal of treatments for HIV-infection^5-9^.

Biochemical and structural studies revealed that slow cleavage of CA-SP1 is due to structural sequestration of the proteolysis site^10-13^. Within the assembled immature HIV-1 Gag lattice, the CA-SP1 junction folds into an α-helix (the junction helix), which self-associates into a 6-helix bundle, stabilizing the Gag hexamer^13,14^. The scissile bond between CA-L231 and SP1-A1 is located in the middle of the junction helix and is occluded inside the 6-helix bundle. Therefore, for the protease to gain access to this site, the 6-helix bundle must at least partially unfold. Although the detailed mechanism of inhibition has not been ascertained, small-molecule MIs, such as 3-*O*-(3’,3’-dimethylsuccinyl)-betulinic acid (Bevirimat or BVM), 1-[2-(4-tert-butylphenyl)-2-(2,3-dihydro-1H-inden-2-ylamino)ethyl]-3-(trifluoromethyl)pyridin-2-one (PF-46396) and their analogs are thought to interfere with proteolysis by binding to the CA-SP1 junction and stabilizing the 6-helix bundle^6,7,15-18^. Thus, MIs do not directly interfere with substrate binding but rather act indirectly by inhibiting the unfolding of the 6-helix bundle and in effect impeding access of the protease to its substrate.

Despite being potent inhibitors of HIV infection in laboratory settings, MIs have not yet been approved for clinical use. BVM underwent phase I and phase II clinical trials, during which significant, dose-dependent viral load reductions in HIV-1-infected individuals were observed^19^. However, further studies revealed that in up to 50% of patients, BVM did not affect viral loads^20,21^. This BVM resistance is associated with naturally occurring viral sequence polymorphs, in particular SP1 amino acid changes at residues 7 and 8 (SP1-V7A, -V7M, -T8Δ and -T8N)^20^. In addition, BVM resistant variants were generated through multiple rounds of selection against BVM *in vitro*, resulting in amino acid changes in SP1 residues 1 and 3 (SP1-A1V, -A3T, and -A3V); of these, SP1-A1V does not impair viral replication^10^.

Inositol hexakisphosphate (IP6), a negatively charged small molecule that is abundant in cells, also stabilizes the CA-SP1 junction by binding to the 6-helix bundle. In contrast to BVM, which binds in the center of the helical bundle^14,18^, IP6 is located just above the 6-helix bundle and forms salt bridges with two rings of lysine side chains (CA-K158 and CA-K227)^22^. Although BVM also contains negatively charged carboxylates, it has not been established whether these can compete with IP6 for interacting with the lysine rings.

To assess how MIs bind to the CA-SP1 site and elucidate the mechanisms that underlie BVM resistance, we determined magic angle spinning (MAS) NMR atomic-resolution structures of microcrystalline complexes of a HIV-1 Gag fragment spanning the CA C-terminal domain (CA_CTD_) and SP1 regions (CA_CTD_-SP1), in the presence of BVM and/or IP6. Structures were calculated based on a large number of distance restraints, which were derived from carbon-carbon and carbon-proton correlations in high-quality spectra. Intermolecular correlations between ligand and protein resonances allowed us to verify simultaneous binding of BVM and IP6, and to unambiguously assign the binding orientation of one BVM molecule inside the CA-SP1 junction 6-helix bundle. Overall, the structures reported herein provide unprecedented atomic-level details of how BVM and IP6 interact with CA_CTD_-SP1, unavailable from any other structural techniques, and explain the structural basis of BVM-mediated maturation inhibition and resistance of SP1-A1V and SP1-V7A variants. Our study also highlights the power of MAS NMR spectroscopy for directly observing and structurally characterizing bound small molecules in large macromolecular assemblies with atomic-level detail.

## RESULTS AND DISCUSSION

### Resonance assignments and distance restraints

Negative-stain transmission electron microscopy (TEM) images confirmed previous findings that CA_CTD_-SP1 formed microcrystalline assemblies in the presence of IP6^22^ and that the assemblies appeared similar in the presence or absence of BVM (Fig. 1a). MAS NMR experiments were conducted using eleven sets of samples, prepared with different combinations of isotopic labels, and in the presence or absence of BVM and/or IP6 (summarized in Supplementary Table 1). A total of fourteen one-dimensional (1D), seventy two-dimensional (2D), and six three-dimensional (3D) spectra were recorded. The sensitivity and the resolution of the data sets are exceptionally high and permitted almost complete (96%) backbone resonance assignments. Overall, 8377 cross peaks were assigned (Table 1; all assignments are summarized in Supplementary Fig. 1 and Supplementary Table 2). Importantly, the MAS NMR spectra provide clear ^13^C chemical shift signatures for mature vs. immature lattices (see Supplementary Fig. 2). All CA_CTD_ tail and SP1 residues, except SP1-M14, give rise to distinct, well-resolved peaks in the MAS NMR spectra (shown in Fig. 1b for a stretch of SP1 residues Q6 through I13). The secondary chemical shifts unequivocally indicate that SP1 residues 1-10 are helical in the absence and presence of BVM. Such helical conformation is fully consistent with the known 6-helix bundle structure of SP1 in the immature Gag lattice^13,14,18,22^, in contrast to the structure in high-salt assembled CA-SP1 without IP6 where SP1 residues are dynamically disordered as described previously^23^.

**Table 1.** Summary of samples and the number of assigned peaks

| ID | Sample | Assigned peak type | No of assigned peak* |
| --- | --- | --- | --- |
| 1. | U-<sup>13</sup>C,<sup>15</sup>N-CA<sub>CTD</sub>-SP1/BVM/IP6 | Intra-residue | 1033 |
| | | Sequential ($\vert i-j\vert = 1$) | 372 |
| | | Medium range ($1 < \vert i-j\vert < 4$) | 718 |
| | | Long range ($\vert i-j\vert \geq 5$) | 735 |
| 2. | U-<sup>13</sup>C,<sup>15</sup>N-CA<sub>CTD</sub>-SP1/ IP6 | Intra-residue | 524 |
| | | Sequential ($\vert i-j\vert = 1$) | 38 |
| 3. | U-<sup>13</sup>C,<sup>15</sup>N-CA<sub>CTD</sub>-SP1/BVM/SO<sub>4</sub> | Intra-residue | 776 |
| | | Sequential ($\vert i-j\vert = 1$) | 394 |
| | | Medium range ($1 < \vert i-j\vert < 4$) | 2 |
| | | Long range ($\vert i-j\vert \geq 5$) | 1 |
| 4. | U-<sup>13</sup>C,<sup>15</sup>N-CA<sub>CTD</sub>-SP1/SO<sub>4</sub> | Intra-residue | 393 |
| 5. | U-<sup>13</sup>C,<sup>15</sup>N,<sup>2</sup>H-CA<sub>CTD</sub>-SP1/BVM/IP6 | Intra-residue | 618 |
| | | Sequential ($\vert i-j\vert = 1$) | 26 |
| | | Medium range ($1 < \vert i-j\vert < 4$) | 3 |
| | | Long range ($\vert i-j\vert \geq 5$) | 14 |
| 6. | U-<sup>13</sup>C,<sup>15</sup>N,<sup>2</sup>H-CA<sub>CTD</sub>-SP1/IP6 (Buffer A) | Intra-residue | 431 |
| | | Sequential ($\vert i-j\vert = 1$) | 2 |
| | | Long range ($\vert i-j\vert \geq 5$) | 5 |
| 7. | U-<sup>13</sup>C,<sup>15</sup>N,<sup>2</sup>H-CA<sub>CTD</sub>-SP1/IP6 (Buffer B) | Intra-residue | 361 |
| | | Sequential ($\vert i-j\vert = 1$) | 33 |
| | | Medium range ($1 < \vert i-j\vert < 4$) | 1 |
| | | Long range ($\vert i-j\vert \geq 5$) | 1 |
| 8. | U-<sup>13</sup>C,<sup>15</sup>N,<sup>2</sup>H- CA<sub>CTD</sub>-SP1-V7A/BVM/IP6 | Intra-residue | 491 |
| | | Sequential ($\vert i-j\vert = 1$) | 8 |
| | | Medium range ($1 < \vert i-j\vert < 4$) | 1 |
| | | Long range ($\vert i-j\vert \geq 5$) | 10 |
| 9. | U-<sup>13</sup>C,<sup>15</sup>N,<sup>2</sup>H-CA<sub>CTD</sub>-SP1-V7A/IP6 | Intra-residue | 457 |
| | | Sequential ($\vert i-j\vert = 1$) | 8 |
| | | Medium range ($1 < \vert i-j\vert < 4$) | 1 |
| | | Long range ($\vert i-j\vert \geq 5$) | 6 |
| 10. | U-<sup>13</sup>C,<sup>15</sup>N,<sup>2</sup>H-CA<sub>CTD</sub>-SP1-A1V/BVM/IP6 | Intra-residue | 445 |
| | | Sequential ($\vert i-j\vert = 1$) | 6 |
| | | Long range ($\vert i-j\vert \geq 5$) | 6 |
| 11. | U-<sup>13</sup>C,<sup>15</sup>N,<sup>2</sup>H-CA<sub>CTD</sub>-SP1-A1V/IP6 | Intra-residue | 445 |
| | | Sequential ($\vert i-j\vert = 1$) | 6 |
| | | Long range ($\vert i-j\vert \geq 5$) | 6 |
| **Total assigned peaks** |  |  | **8377** |

\*Cross peaks present in different experiments are counted only once.

**Figure 1.**
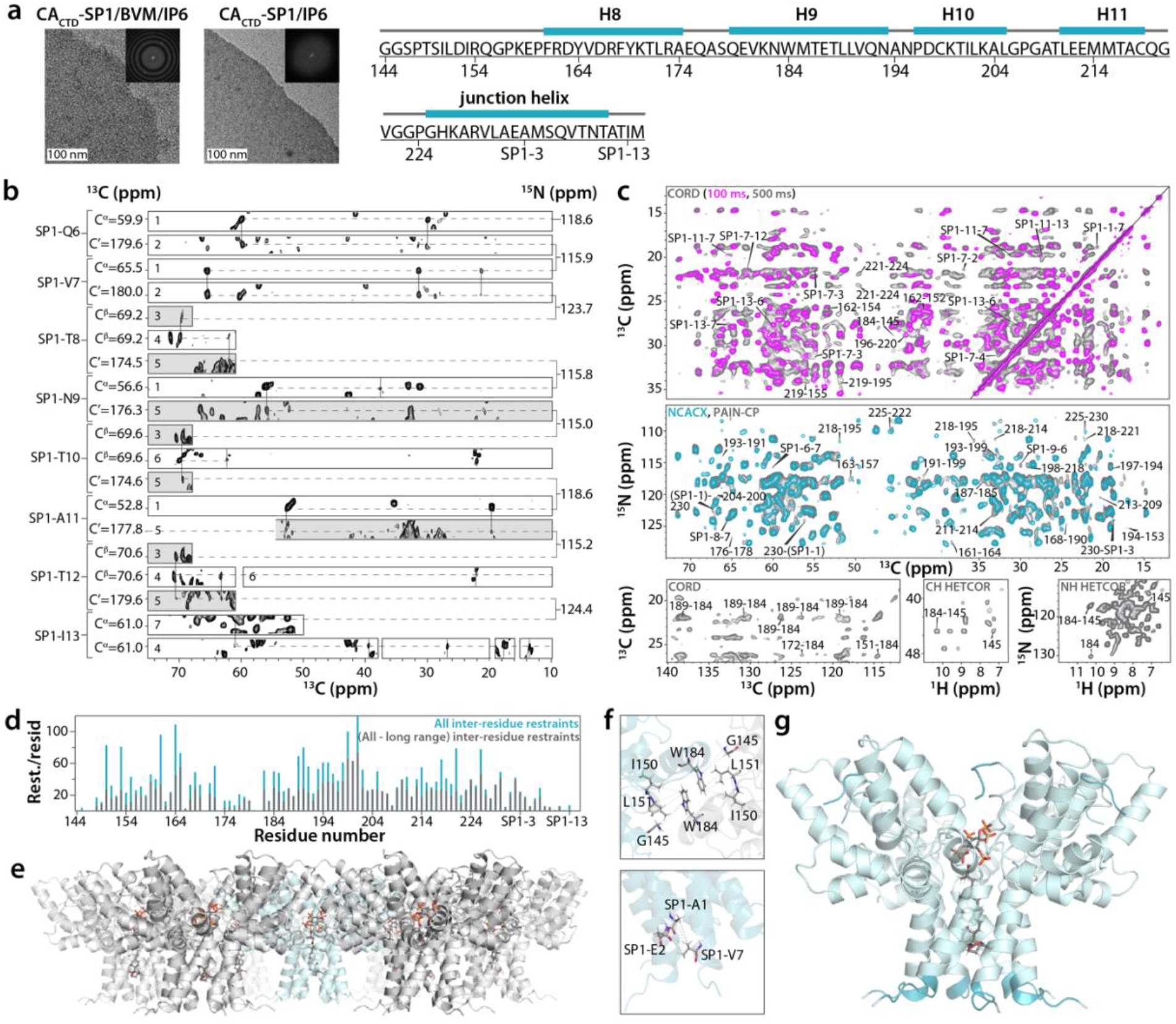
MAS NMR spectra and structure of CA_CTD_-SP1 crystalline array. **a)** Negative stain TEM images of CA_CTD_-SP1 microcrystals assembled with IP6, in the presence (left) or absence of BVM (middle). Insets show the computed Fourier transforms of the images, indicating the expected hexagonal lattices and unit cell spacings. The scale bars are 100 nm. Amino acid sequence of CA_CTD_-SP1 (right). **b)** Representative strips of 3D and 2D MAS NMR spectra of U-^13^C,^15^N-CA_CTD_-SP1/BVM/SO_4_ (white strips) and U-^13^C,^15^N-CA_CTD_-SP1/IP6 (gray strips) crystalline arrays, illustrating sequential assignments for SP1 residues Q6-I13. The MAS NMR spectra are labeled as follows: 3D NCACX (1), 3D NCOCX (2), 2D NCACX at -79 °C (3), 2D INADEQUATE (4), 2D NCOCX at -79 °C (5), 2D CORD (6), 2D NCACX (7). No significant chemical shift perturbations were detected for the free vs. BVM- or IP6-bound samples (see Supplementary Fig. 3). **c)** Top panel: Superposition of selected regions of 2D CORD spectra of U-^13^C,^15^N-CA_CTD_-SP1/BVM/IP6 crystalline arrays for different mixing times: 100 ms (magenta) and 500 ms (gray). Middle panel: Superposition of selected regions of 2D NCACX (cyan) and 2D PAIN-CP (gray) spectra of U-^13^C,^15^N-CA_CTD_-SP1/BVM/IP6 crystalline arrays. Unambiguous long-range and inter-residue correlations of SP1 residues are labeled by amino acid number in the sequence. Bottom panel: Inter-hexamer correlations are shown for the selected regions of 2D CORD (left) and CH HETCOR (middle) spectra of U-^13^C,^15^N-CA_CTD_-SP1/BVM/IP6, and NH HETCOR (right) spectra of U-^13^C,^15^N,^2^H-CA_CTD_-SP1/IP6 (Buffer B). **d)** Number of long-range and all inter-residue MAS NMR restraints per residue plotted against the residue number. **e)** Side view of hexamer of hexamers of BVM- and IP6-bound CA_CTD_-SP1 arrays. **f)** Expansion of inter-hexamer (top panel) and inter-chain (bottom panel) regions showing distance restraints obtained from MAS NMR correlation experiments. **g)** MAS NMR structure of a single hexamer of BVM and IP6-bound CA_CTD_-SP1 crystalline array. The residues detected by MAS NMR and not modeled in the X-ray and cryo-EM structures^13,14^ are shown in darker cyan.

A large number of intra-protein correlations were detected in multiple MAS NMR spectra (Fig. 1c), and, given the excellent resolution, no ^13^C isotopically diluted samples were needed to distinguish intramolecular from intermolecular correlations: all cross peaks are well resolved and could be unambiguously assigned, permitting to extract distance restraints. In total, 3048 non-redundant unambiguous protein-protein distance restraints (^13^C-^13^C, ^15^N-^13^C, ^13^C-^1^H, and ^15^N-^1^H) were obtained, comprising 674 medium-range (1<|i-j|<4), 627 long-range (|i-j|≥5), with 39 long-range inter-chain and 22 long-range inter-hexamer restraints. With nearly 30 non-redundant unambiguous restraints per residue or over 52% C-C restraint completeness (see Table 2) and a very large number of inter-residue restraints (1754) derived from long-range correlations (Fig. 1d), this study yielded the highest number of distance restraints of any protein MAS NMR investigation to date. Correlations were seen between CA-K158Cε and IP6-H2/H4/H6, indicating that CA-K158 mediates IP6 binding. They were translated into 6 restraints, involving two adjacent chains in the 6-helix bundle for each of the three IP6 protons. Additionally, a very weak correlation was observed with CA-K227Cδ. Correlations with BVM resulted in 7 distance restraints, which unambiguously defined the binding orientation of the inhibitor within the 6-helix bundle. These correlations are summarized in Fig. 2a. A summary of all distance restraints is provided in Table 2 and a plot of all inter-residue contacts is illustrated in Supplementary Fig. 4.

**Table 2.**
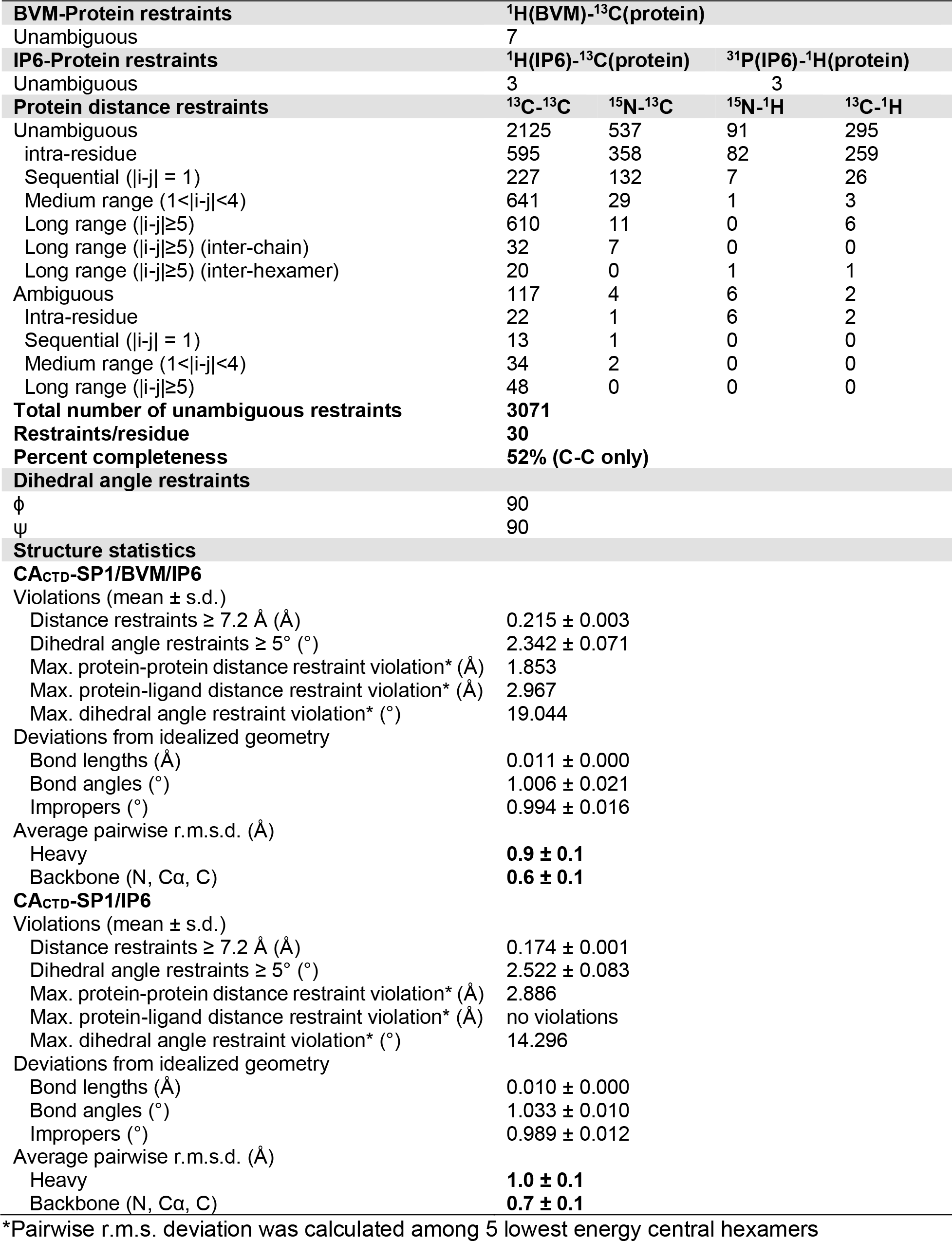
Summary of MAS NMR restraints and structure statistics

**Figure 2.**
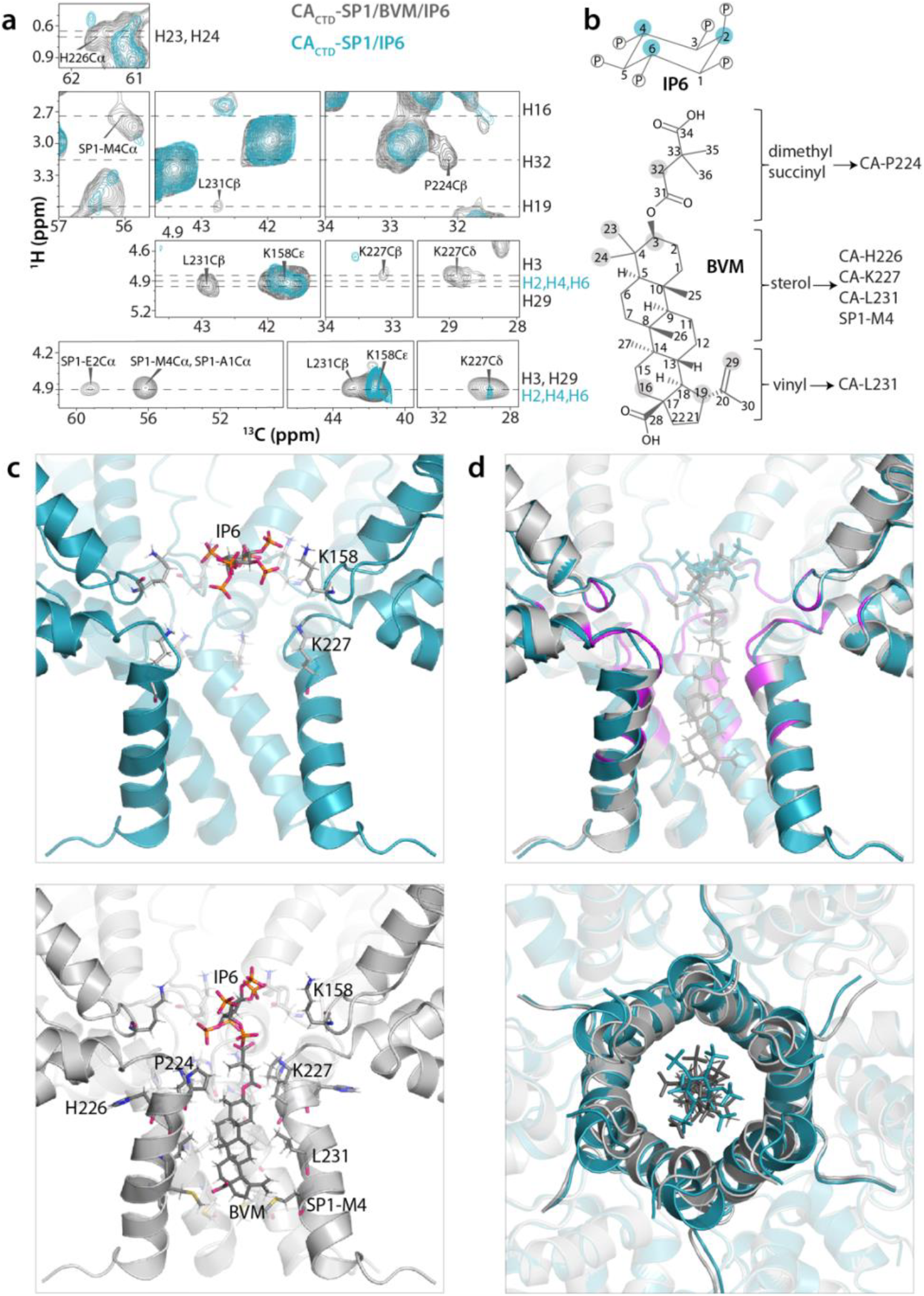
MAS NMR structure of BVM- and IP6-bound CA_CTD_-SP1. **a)** Superposition of selected regions of 2D HC CP HETCOR spectra (top panel) and dREDOR-HETCOR spectra (bottom panel) of U-^13^C,^15^N,^2^H-CA_CTD_-SP1/BVM/IP6 (grey) and U-^13^C,^15^N,^2^H-CA_CTD_-SP1/IP6 (cyan) assemblies. The BVM and IP6 ^1^H atoms and CA_CTD_-SP1 residues with ^13^C atoms are indicated outside and inside each spectrum, respectively. **b)** Chemical identity of IP6 and BVM molecules. The IP6 and BVM protons interacting with CA_CTD_-SP1 residues are shown in cyan and gray, respectively. **c)** Top panel: IP6 binding mode in the hexamer of CA_CTD_-SP1/IP6 assemblies (PDB 7R7Q, this work). Bottom panel: IP6 and BVM binding modes in the hexamer of CA_CTD_-SP1/BVM/IP6 assemblies (PDB 7R7P, this work). Residues interacting with IP6 or BVM are shown as sticks. **d)** Superposition of MAS NMR structure of CA_CTD_-SP1/BVM/IP6 and CA_CTD_-SP1/IP6 shown from side view (top) and top view (bottom). BVM binding induces major structural rearrangements of the SP1 helices, resulting in the tightening of the pore and quenching the motions of the simultaneously bound IP6. Residues colored in magenta give rise to high intensity peaks corresponding to intra- and inter-residue correlations upon BVM binding.

### Higher-order structure of CA_CTD_-SP1 and conformations of BVM and IP6

Structures (Fig. 1e) were calculated by integrating the experimental MAS NMR restraints (i.e., protein-protein, protein-BVM, and protein-IP6 distance and protein torsion angle restraints) and a hexamer-of-hexamers structural envelope generated from X-ray coordinates of the CA_CTD_-SP1 hexamer (PDB 5I4T)^13^. The hexamer-of-hexamers unit represents the minimal building block that recapitulates the critical inter-hexamer interfaces in the immature capsid lattice. The CA_CTD_-SP1 MAS NMR-derived inter-chain and inter-hexamer contacts are shown in Fig. 1f.

As described previously^13,14,18,22^, the CA_CTD_-SP1 hexamer exhibits the shape of a goblet, with the globular CA_CTD_ domain forming the cup and the 6-helix bundle CA-SP1 junction fashioning the stem. In the crystal structure of CA_CTD_-SP1 and the cryo-EM structure of full-length Gag, the junction helix terminates around residue SP1-V7, with the remaining residues not visible, likely due to conformational disorder^13,18,22^. Remarkably, the entire SP1 region, including the SP1 tail, except for SP1-M14, is well defined in the MAS NMR structures (Fig. 1g and Supplementary Fig. 5 and 6). MAS NMR chemical shifts predict a helical conformation up to SP1-T10, with the last 4 residues (A11-M14) being in a random coil structure (Fig. 1g). For SP1-E2, S5, T8, N9, T10, T12, and I13 residues, peak intensities are low, suggesting conformational heterogeneity (Fig. 1b), consistent with the X-ray^13^ and cryo-EM data^14^.

In the BVM-bound structure, the inhibitor is located inside the channel formed by the 6-helix bundle, with the sterol ring occupying the hydrophobic interior of the channel and making contacts with protein residues CA-H226, K227, L231, and SP1-M4. The dimethyl succinyl moiety is oriented towards the CA_CTD_ goblet, as judged by H32(BVM)-P224Cβ(protein) and H3(BVM)-K227Cβ(protein) correlations, whereas the vinyl group points towards the SP1 tail, as suggested by H19(BVM)-L231Cβ(protein) and H29(BVM)-L231Cβ(protein) correlations (Fig. 2a,b). In summary, our structure confirms that BVM organizes the CA-SP1 junction by binding inside the central channel of the 6-helix bundle^14,18^, and, importantly, defines the orientation of the bound drug.

Comparison of the structures in the presence and absence of BVM clearly shows that BVM binding results in apparent tightening of the hexamer pore (Fig. 2c,d). We also observed pronounced conformational heterogeneity of CA-P157, K158, E159, and SP1-M4, which indicates asymmetry of the six protein side chain copies in the hexameric ring. This result provides further evidence that a single, non-symmetric BVM molecule is bound inside the 6-helix bundle (Supplementary Fig. 7). Additionally, the SP1 tail becomes less dynamic upon BVM binding, as evidenced by increased intensity for cross peaks of tail residues, with a concomitant reorientation of side chains in the structure close to the binding site, such as CA-P157, K158, E159, N195, G220, V221, G222, G223, P224, K227, V230, L231, and SP1-E2 (Fig 2d and 3a, top panels, and Supplementary Fig. 8). We suggest that these structural changes all contribute to the stabilizing effect of BVM binding.

**Figure 3.**
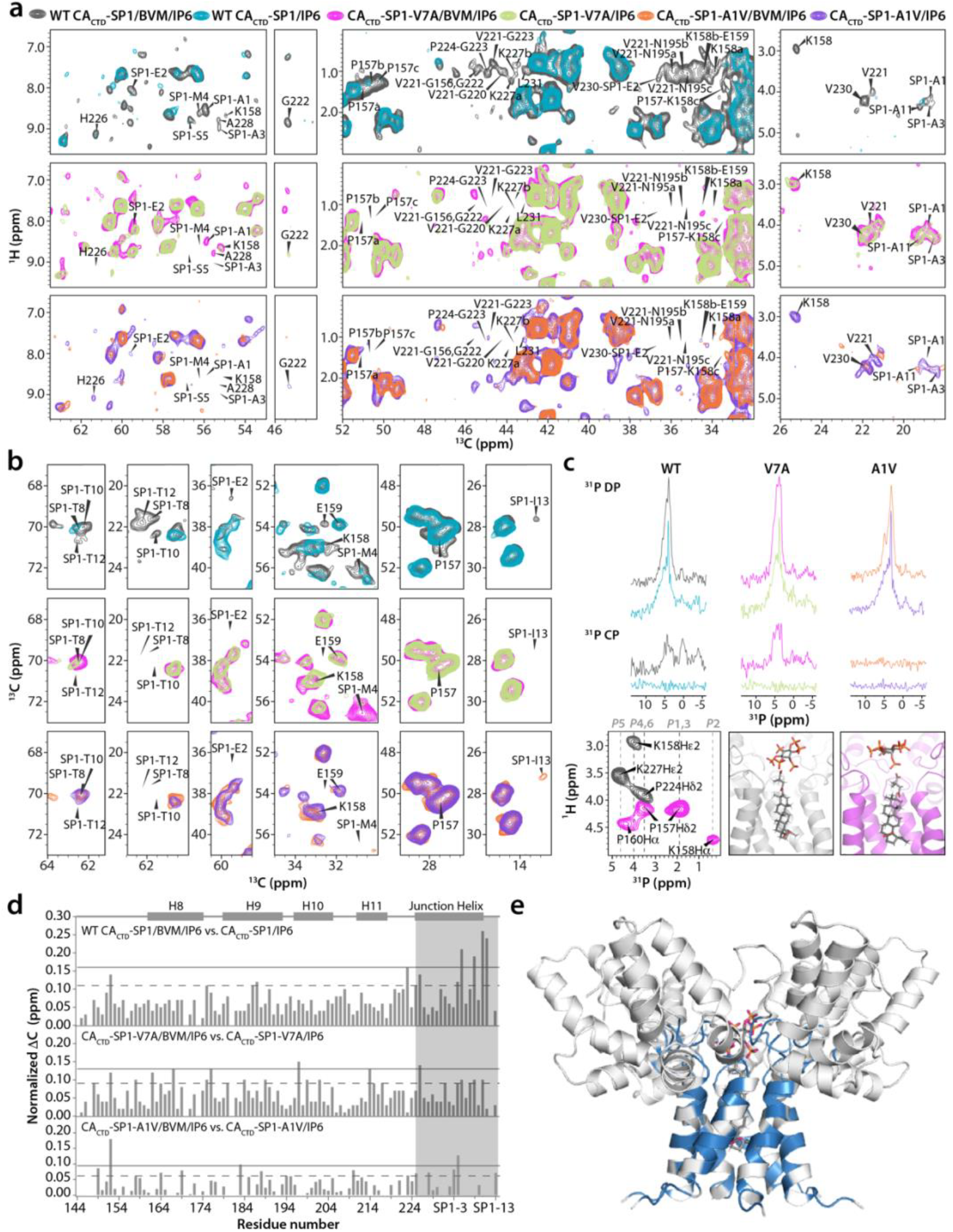
Effect of BVM binding on wild-type (WT) and the SP1-A1V and SP1-V7A variants. **a)** Superposition of selected regions of 2D HC CP HETCOR spectra of U-^13^C,^15^N,^2^H-CA_CTD_-SP1/BVM/IP6 (gray) and U-^13^C,^15^N,^2^H-CA_CTD_-SP1/IP6 (cyan) (top panel); U-^13^C,^15^N,^2^H-CA_CTD_-SP1-V7A/BVM/IP6 (magenta) and U-^13^C,^15^N,^2^H-CA_CTD_-SP1-V7A/IP6 (light green) (middle panel); and U-^13^C,^15^N,^2^H-CA_CTD_-SP1-A1V/BVM/IP6 (orange) and U-^13^C,^15^N,^2^H-CA_CTD_-SP1-A1V/IP6 (purple) (bottom panel). Intra- and inter-residue correlations arising upon BVM binding only in WT CA_CTD_-SP1 but not in CA_CTD_-SP1-V7A or CA_CTD_-SP1-A1V are labeled. Residues showing multiple conformers are denoted with a,b,c. **b)** Superposition of selected regions of 2D CORD spectra of U-^13^C,^15^N,^2^H-CA_CTD_-SP1/BVM/IP6 (gray) and U-^13^C,^15^N,^2^H-CA_CTD_-SP1/IP6 (cyan) (top panel); U-^13^C,^15^N,^2^H-CA_CTD_-SP1-V7A/BVM/IP6 (magenta) and U-^13^C,^15^N,^2^H-CA_CTD_-SP1-V7A/IP6 (light green) (middle panel); and U-^13^C,^15^N,^2^H-CA_CTD_-SP1-A1V/BVM/IP6 (orange) and U-^13^C,^15^N,^2^H-CA_CTD_-SP1-A1V/IP6 (purple) (bottom panel). Intra-residue cross-peaks exhibiting intensity or chemical shift changes due to BVM binding in WT CA_CTD_-SP1 are labeled. The corresponding correlations are either absent or exhibit small chemical shift perturbations upon BVM binding to CA_CTD_-SP1-V7A/BVM/IP6 and in CA_CTD_-SP1-A1V/BVM/IP6. **c)** Top: ^31^P direct polarization (DP, top) and cross polarization (CP, middle) spectra of U-^13^C,^15^N,^2^H-CA_CTD_-SP1/BVM/IP6 (gray), U-^13^C,^15^N,^2^H-CA_CTD_-SP1/IP6 (cyan), U-^13^C,^15^N,^2^H-CA_CTD_-SP1-V7A/BVM/IP6 (magenta), U-^13^C,^15^N,^2^H-CA_CTD_-SP1-V7A/IP6 (light green), U-^13^C,^15^N,^2^H-CA_CTD_-SP1-A1V/BVM/IP6 (orange), and U-^13^C,^15^N,^2^H-CA_CTD_-SP1-A1V/IP6 (purple). Bottom: Overlay of 2D (H)PH HETCOR spectra of U-^13^C,^15^N,^2^H-CA_CTD_-SP1/BVM/IP6 (gray) and U-^13^C,^15^N,^2^H-CA_CTD_-SP1-V7A/BVM/IP6 (magenta) (left). Tilted orientation of IP6 in WT CA_CTD_-SP1/BVM/IP6 (middle) (PDB 7R7P, this work). Horizontal orientation of IP6 in CA_CTD_-SP1-V7A/BVM/IP6 (right). **d)** Chemical shift perturbations induced by BVM binding for WT CA_CTD_-SP1 (top), CA_CTD_-SP1-V7A (middle), and CA_CTD_-SP1-A1V (bottom), plotted against residue number. **e)** CA_CTD_-SP1/BVM/IP6 structure with residues exhibiting unique CSPs, intra- and inter-residue correlations, and enhanced peak intensities upon BVM binding to WT CA_CTD_-SP1 shown in blue.

Importantly, our data revealed that BVM and IP6 can bind to the CA_CTD_-SP1 hexamer simultaneously, as proposed recently^24,25^. This is supported by the observation of distinct sets of protein-BVM and protein-IP6 cross peaks in the same spectra. Importantly, the mode of interaction between IP6 and CA_CTD_-SP1 is distinct in the presence and absence of BVM. Specifically, the ^1^H-^13^C HETCOR and dREDOR-HETCOR data sets reveal correlations between multiple protons of BVM (H3, H16, H19, H23, H24, H29, H32) and IP6 (H2, H4, H6) with different CA_CTD_-SP1 residues (Fig. 2a), implying that IP6 and BVM binding is not competitive. In the absence of BVM, IP6 binds horizontally inside the neck region of the channel inside the 6 -helix bundle, coordinated by six CA-K158 residues. This configuration is supported by correlations between the IP6 H2/H4/H6 group of resonances (chemical shifts are too close to be assigned to individual atoms) and the Cε atoms of six CA-K158 sidechains (Fig. 2a). A weak H2/H4/H6(IP6) correlation to CA-K227Cd is also observed, confirming a specific contact between IP6 and this residue, consistent with recent reports^22^. Interestingly, no equivalent correlations are observed in the ^1^H-^31^P 1D CPMAS or 2D HETCOR spectra (Fig. 3c), indicating that coordination by the six lysine residues is dynamically averaged, with IP6 undergoing local motions inside the pore. In contrast, when BVM is bound, three intense cross peaks appear in the ^1^H-^31^P 2D HETCOR spectrum of U-^13^C,^15^N,^2^H-CA_CTD_-SP1/BVM/IP6 (Fig. 3c), corresponding to correlations between 3 different phosphorus atoms of IP6 and the sidechains of CA-K158, K227, and P224 residues. The presence of intense cross peaks implies that the motion of IP6 in the presence of BVM is arrested. Whether the arrest of IP6 motion is connected to the stabilization the 6-helix bundle is unclear at present, although bound IP6 dynamics can be used to assess MI activity (see below).

### BVM binding to SP1-A1V and SP1-V7A variants

Understanding the mechanisms of resistance to MI inhibitors is of key importance for further development of such molecules as drugs. We therefore evaluated how CA_CTD_-SP1 variants associated with BVM resistance affect binding. CA_CTD_-SP1 assemblies harboring the SP1-A1V and V7A substitutions were prepared and 2D ^1^H-^13^C HETCOR, ^13^C-^13^C CORD, 1D ^31^P DP and CPMAS, and 2D ^1^H-^31^P HETCOR spectra were recorded in the presence and absence of BVM and/or IP6 (Fig. 3a-c). In the absence of BVM, the peak intensities for most SP1 residues in the A1V and V7A variants are considerably lower than for the wild-type (WT) protein, and many correlations associated with other regions are missing (Supplementary Fig. 9). This indicates that in both variants, the SP1 regions are inherently more dynamic than in the WT protein. Comparison of ^13^C chemical shift perturbations (CSPs) induced by BVM for the WT and the two variants, A1V and V7A, revealed most extensive CSPs for the WT assemblies, especially for resonances associated with residues in SP1 (M4, S5, T8, T10, A11), the type-II β-turn (CA-G223, G225) and the junction helix (CA-H226) (Fig. 3d,e). For A1V variant, we found none of the protein-BVM correlations that were observed with WT CA_CTD_-SP1, indicating that BVM binds very weakly, is very mobile or does not bind at all. In contrast, we observed modest, but statistically significant CSPs with V7A, indicating that this variant bound to BVM more efficiently than A1V (but less efficiently than WT). Interestingly, BVM binding to the V7A assemblies did not render the complex more rigid, in contrast to observations for WT CA_CTD_-SP1, since none of the equivalent correlations are present in the MAS NMR spectra (Fig. 3a, and 3b). Consistent with the CSP data, ^31^P signals of IP6 were observed in 1D ^31^P CPMAS spectra of CA_CTD_-SP1-V7A/BVM/IP6 assemblies, (Fig. 3c), similar to findings for WT CA_CTD_-SP1/BVM/IP6, while they are absent in CA_CTD_-SP1-A1V/BVM/IP6 samples. Therefore, we conclude that IP6 is motionally restricted in the presence of BVM in CA_CTD_-SP1-V7A assemblies, but mobile in assemblies of CA_CTD_-SP1-A1V. Our data correlate well with observations from virology studies^26,27^ that showed that the virions harboring the SP1-A1V mutation bind BVM less efficiently than WT in virus-like particles (VLPs)^27^. In addition, the CA-SP1 processing is faster in these mutant virions, compared to WT virus, consistent with the notion that SP1-A1V variant possesses a more conformationally dynamic 6-helix bundle^10^. In contrast, CA-SP1 processing in SP1-V7A and WT virions is similar^21^ suggesting different BVM resistance mechanisms for the SP1-V7A and SP1-A1V sequence variants^26^. Taken together, our MAS NMR results suggest that distinct mechanisms can be associated with loss of BVM binding to MI resistant variants.

2D ^1^H-^31^P HETCOR spectra of WT CA_CTD_-SP1/BVM/IP6 and CA_CTD_-SP1-V7A/BVM/IP6 assemblies reveal distinct ^31^P(IP6)-^1^H(protein) cross peaks (Fig. 3c). For the WT protein, three correlations are present: P4/6(IP6)-K158Hε2(protein), P5(IP6)-K227Hε2(protein), and P4/6(IP6)-P224Hδ2(protein). For CA_CTD_-SP1-V7A, P1/3/4/6(IP6)-P157Hδ2(protein), P2(IP6)-K158Hα(protein), and P4/6(IP6)-P160Hα(protein) correlations are observed (Fig. 3c and Supplementary Fig. 10 and 11). In contrast, no correlations are seen in the CA_CTD_-SP1-A1V spectra, suggesting that IP6 is dynamic. For WT CA_CTD_-SP1, the correlation between a single phosphorus atom (P5) of IP6 and SP1-K227Hε2 suggests that IP6 is facing only one of the six CA-K227 residues. The fact that no correlations to P2 and P1,3 are seen in WT CA_CTD_-SP1 spectra is consistent with a tilted orientation of IP6 (Fig. 3c). For the SP1-V7A variant, the absence of IP6 correlations with CA-K227, together with the presence of IP6 correlations with CA-P157, K158, and P160, suggests that IP6 adopts a horizontal orientation in the neck region (Fig. 3c). We posit that the distinct dynamic properties and orientations adopted by IP6 in the WT, SP1-A1V, and SP1-V7A protein assemblies may correlate with BVM inhibitory activity against the resistant viruses, suggesting the tantalizing possibility that ^31^P MAS NMR of IP6 bound assemblies could be developed for MI screening.

## CONCLUSIONS

Here, we presented MAS-NMR structures of the CA_CTD_-SP1 lattice in complex with IP6, in the presence or absence of BVM that reveal novel atomic-level details of protein-ligand interactions. In particular, our data confirm that BVM binds inside the pore of the CA-SP1 6-helix bundle, simultaneously with IP6. Importantly, we unambiguously defined the binding orientation of both ligands and showed that BVM causes pore tightening associated with structural rearrangements of residues in the SP1 helices and quenches the dynamics of the simultaneously bound IP6, consistent with a stabilization of the 6-helix bundle^14-16,18,28^. Additionally, we uncovered previously unknown effects of BVM on IP6 dynamics that likely contribute to preventing protease access to the CA-SP1 cleavage site. Furthermore, we discovered that BVM resistance in SP1-A1V and SP1-V7A variants is associated with loss of MI binding or loss of a stable 6-helix bundle conformation, respectively.

In addition to providing important molecular insights into BVM-mediated effects for Gag maturation, our study also represents a major technological advance in MAS NMR, beyond commonly used approaches for characterization of bound ligands requiring their ^13^C isotopic labeling. This was possible through judicious exploitation of ^1^H resonances of BVM as well as ^1^H and ^31^P resonances of IP6. Moreover, deuteration of CA_CTD_-SP1 and dREDOR-filtered experiments proved critical to this end. To the best of our knowledge, the present MAS NMR structures of higher-order assemblies are the first of this kind without necessitating ^13^C isotopic dilution.

Taken together, the findings presented here not only elucidate the atomic details of BVM binding to HIV-1 CA_CTD_-SP1 and inform on BVM-resistance mechanisms, but also may suggest new strategies for the design of more-potent next-generation MIs.

## MATERIALS AND METHODS

### Protein expression and purification

The expression plasmid for the CA_CTD_-SP1 fragment of HIV-1 strain NL4-3 Gag containing a non-cleavable His-tag and mutation P373T (SP1-P10T) was described previously^13^. The SP1-V7A and A1V mutations were added to the construct by Quikchange mutagenesis (Agilent). The SP1-T10 sequence polymorph exhibits the same infectivity and BVM antiviral activity (IC_50_) profiles as the SP1-P10, as shown in Supplementary Fig. 12. Proteins were expressed in transformed E. coli BL21 (DE3) cells, which were grown in a shaker incubator at 37 °C until mid-log phase (OD_600_ of 0.6-1) and then induced with 1 mM IPTG overnight at 18 °C. Expression of U-^13^C,^15^N enriched and U-^13^C,^15^N,^2^H enriched CA_CTD_-SP1 proteins had additional pre-culturing steps to slowly acclimate the cells from rich media to minimal media, and performed as reported previously^29,30^. Cells were harvested by centrifugation and stored at -80 °C until use.

Protein purification was performed as previously described^13^. In brief, bacterial pellets were resuspended in 50 mM Tris, pH 8.3, 1 M LiCl, containing protease inhibitor tablets (Roche) and supplemented with 0.3% (w/v) deoxycholate. Cells were lysed by incubation with lysozyme, followed by sonication. Lysates were clarified by centrifugation, filtered, and incubated with Ni-NTA agarose resin (Qiagen) for 30 min at 4 °C. Unbound fractions were washed away, and bound protein was eluted by a step gradient of 15 mM imidazole to 300 mM imidazole. Protein was purified to homogeneity using anion exchange chromatography in 20 mM Tris, pH 8.0, 0 .5 M NaCl. Pure protein was concentrated to 10 mg/mL, flash-frozen in liquid nitrogen, and stored at - 80 °C until use.

### Buffer exchange

Two sets of U-^13^C,^15^N,^2^H-CA_CTD_-SP1/IP6 samples were prepared, one in Buffer A (20 mM Tris, pD 8.0, 0.5 M NaCl; made from 1 M Tris stock at 99.9% purity in D_2_O (Cambridge) and pD adjusted with deuterium chloride (Sigma) prepared with D_2_O at 99.9% purity (Cambridge)) and the second in Buffer B (20 mM Tris, pH 8.0, 0.5 M NaCl). The protein samples in Buffer A were prepared by buffer exchange as follows: 0.5 mL protein at 10 mg/mL was diluted into 10 mL with Buffer A then re-concentrated via centrifugation, 4 times. After exchange, the samples were recovered in a final volume 0.5 mL, with concentrations between 9-10 mg/mL.

### Protein assembly and sample preparation

Proteins were assembled with 1.6 mM IP6 (Sigma-Aldrich) and 1.4 mM BVM (Sigma-Aldrich), for a final reaction volume of 1 mL. Some samples that were used to obtain or confirm resonance assignments were assembled by mixing protein with equal volume of 1.5 M ammonium sulfate (Sigma-Aldrich), 0.1 M Tris, pH 8.5. Assemblies were incubated overnight. Optimal assemblies were obtained at 16-20 °C incubation temperatures. U-^13^C,^15^N labeled assemblies (∼50 mg protein) were centrifuged at 10,000x*g* and packed in 3.2 mm Bruker thin-walled rotors. Buffer exchanged protein assemblies (∼17 mg protein) were centrifuged at 10,000x*g* and packed in 1.9 mm Bruker rotors.

### BVM antiviral activity and particle infectivity

The activity of BVM against SP1-P10 and the SP1-T10 derivative was determined essentially as reported previously^26^. Briefly, HEK 293T cells were transfected with pNL4-3/SP1-P10 or pNL4-3/SP1-T10 molecular clones, and the transfected cells were treated with 24 concentrations of BVM ranging from 0 to 10 μM. Virus-containing supernatants were harvested, normalized for reverse transcriptase (RT) activity, and used to infect the TZM-bl indicator cell line. Infectivity data were analyzed with GraphPad Prism 7 for Mac OS X from three independent experiments. Curves were fit using nonlinear regression as log(inhibitor) versus normalized response, with a variable slope using a least-squares (ordinary) fit.

To measure the relative infectivity of SP1-P10 vs. the SP1-T10 derivative, HEK 293T cells were transfected with the pNL4-3/SP1-P10 or pNL4-3/SP1-T10 molecular clones. Virus-containing supernatants were harvested, normalized for RT activity, and used to infect the TZM-bl indicator cell line. Luciferase activity was measured at 2 days postinfection. The specific infectivities are presented relative to those of the SP1-P10 (100%). Error bars indicate standard deviations (n = 4 independent assays performed in duplicate).

### MAS NMR spectroscopy

MAS NMR experiments of U-^13^C,^15^N-CA_CTD_-SP1/BVM/IP6, U-^13^C,^15^N-CA_CTD_-SP1/IP6, U-^13^C,^15^N-CA_CTD_-SP1/BVM/SO_4_, U-^13^C,^15^N-CA_CTD_-SP1/SO_4_ crystalline arrays were performed on Bruker 19.97 T narrow bore AVIII spectrometer outfitted with 3.2 mm E-Free HCN probes. The MAS frequency was 14 kHz controlled to within ± 10 Hz by a Bruker MAS controller. The actual sample temperature was maintained at 4 ± 1 °C throughout the experiments and at -10 ± 1 °C for some specific experiments using the Bruker temperature controller. The Larmor frequencies were 850.4 MHz (^1^H), 213.9 MHz (^13^C), and 86.2 MHz (^15^N) at 19.97 T spectrometer. The typical 90° pulse lengths were 2.6-3.0 μs for ^1^H and 4.3-4.5 μs for ^13^C, and 4.2-4.7 μs for ^15^N. The ^1^H-^13^C and ^1^H-^15^N cross-polarization employed a linear amplitude ramp of 90-110% on ^1^H, and the center of the ramp matched to Hartmann-Hahn conditions at the first spinning sideband, with contact times of 0.7-1.5 ms and 1.0-1.7 ms, respectively. Different 2D combined R2_n_^v^-driven (CORD)^31^ mixing times such as 10 ms, 50 ms, 100 ms, were applied for different experiments and ^1^H field strength during CORD was 14 kHz. Band-selective magnetization transfer from ^15^N to^13^C contact time was 6.0-6.5 ms. SPINAL-64^32^ decoupling (83-95 kHz) was used during the evolution and acquisition periods.

MAS NMR experiments were also performed on Magnex/Bruker 14.1 T AVIII spectrometer outfitted with 3.2 mm E-Free HCN probe. The Larmor frequencies were 599.8 MHz (^1^H), 150.8 MHz (^13^C), and 60.7 MHz (^15^N) at 14.09 T spectrometer. The typical 90° pulse lengths were 2.6-3.0 μs for ^1^H and 4.0-4.7 μs for ^13^C, and 4.2-4.6 μs for ^15^N. The ^1^H-^13^C and ^1^H-^15^N cross-polarization employed a linear amplitude ramp of 90-110% on ^1^H, and the center of the ramp matched to Hartmann-Hahn conditions at the first spinning sideband, with contact times of 1.1-2.0 ms and 1.3-1.8 ms, respectively. Different CORD mixing times such as 25 ms, 100 ms, 250 ms, 500 ms were applied for different experiments and ^1^H field strength during CORD was 14 kHz. Band-selective ^15^N-^13^C SPECIFIC-CP contact time was 4.0-6.0 ms. SPINAL-64^32^ decoupling (80-86 kHz) was used during the evolution and acquisition periods. 2D phase-shifted ^13^C-detected proton-assisted insensitive-nuclei cross polarization (PAIN-CP)^30^ experiment was acquired for U-^13^C,^15^N-CA_CTD_-SP1/BVM/IP6 crystalline array on 14.09 T spectrometer. During the PAIN-CP mixing period, RF field strengths for ^1^H, ^15^N and ^13^C channels were all 60 kHz. The length of the PAIN-CP mixing period was 4 ms.

Low temperature MAS NMR experiments of U-^13^C,^15^N-CA_CTD_-SP1/IP6 were performed on 17.6 T Brucker AVIII spectrometer outfitted with 3.2 mm low T E-Free HCN probe. The MAS frequency was 15kHz controlled to within ± 10 Hz by a Bruker MAS controller. The actual sample temperature was maintained at -37 ± 1 ^0^C throughout the experiments. The Larmor frequencies were 750.1 MHz (^1^H), 188.6 MHz (^13^C), and 76.0 MHz (^15^N) at 17.6 T spectrometer. The typical 90^0^ pulse lengths were 2.5 μs for ^1^H, 3.1 μs for ^13^C, and 3.3 μs for ^15^N. The ^1^H-^13^C and ^1^H-^15^N cross-polarization employed a linear amplitude ramp of 70-100% on ^1^H, and the center of the ramp matched to Hartmann-Hahn conditions at the first spinning sideband, with contact times of 0.6 ms and 0.9 ms, respectively. CORD mixing time of 25 ms was applied for different experiments and ^1^H field strength during CORD was 15 kHz. Band-selective magnetization transfer from ^15^N to ^13^C contact time was 5.0-6.0 ms. SWFTPPM decoupling (80 kHz) was used during the evolution and acquisition periods.

MAS NMR experiments of U-^13^C,^15^N,^2^H-CA_CTD_-SP1/BVM/IP6, U-^13^C,^15^N,^2^H-CA_CTD_-SP1/IP6 crystalline arrays were performed with a 1.9 mm HCN probe on Bruker 19.97 T narrow bore AVIII spectrometer. The typical 90° pulse lengths were 3.0-3.1 μs for ^1^H, 3.9-4.0 μs for ^13^C, and 3.9-4.0 μs for ^15^N. The ^1^H-^13^C and ^1^H-^15^N cross-polarization employed a linear amplitude ramp of 90-110% on ^1^H, and the center of the ramp matched to Hartmann-Hahn conditions at the first spinning sideband, with contact times of 0.3-2.6 ms and 1.4-1.8 ms respectively. The MAS frequency was 40 kHz controlled to within ±10 Hz by a Bruker MAS controller. The actual sample temperature was maintained at 4 ± 1 °C throughout the experiments otherwise specified using the Bruker temperature controller. MAS NMR double-REDOR filtered experiments employed simultaneous ^1^H^13^C/^1^H^15^N REDOR dephasing periods of 5 ms, to eliminate signals from ^1^H directly bonded to ^13^C and ^15^N. The ^1^H-^13^C cross-polarization employed a linear amplitude ramp of 90-110% on ^1^H, and the center of the ramp matched to Hartmann-Hahn conditions at the first spinning sideband, with contact times of 10 ms.

MAS NMR experiments of U-^13^C,^15^N,^2^H-CA_CTD_-SP1/BVM/IP6, U-^13^C,^15^N,^2^H-CA_CTD_-SP1/IP6, U-^13^C,^15^N,^2^H-CA_CTD_-SP1-V7A/BVM/IP6, U-^13^C,^15^N,^2^H-CA_CTD_-SP1-V7A/IP6, U-^13^C,^15^N,^2^H-CA_CTD_-SP1-A1V/BVM/IP6, U-^13^C,^15^N,^2^H-CA_CTD_-SP1-A1V/IP6 crystalline arrays were performed with a 1.9 mm HX probe on Bruker 19.97 T narrow bore AVIII spectrometer. The typical 90° pulse lengths were 2.3-3.1 μs for ^1^H and 3.0-3.3 μs for ^13^C. The ^1^H-^13^C cross-polarization employed a linear amplitude ramp of 90-110% on ^1^H, and the center of the ramp matched to Hartmann-Hahn conditions at the first spinning sideband, with contact times of 2 ms. The MAS frequency was 40 kHz and 14 kHz controlled to within ± 10 Hz by a Bruker MAS controller. The actual sample temperature was maintained at 4 ± 1 °C throughout the experiments otherwise specified using the Bruker temperature controller.

^31^P solid state MAS NMR spectra were acquired with a 1.9 mm HX probe on 19.97 T Bruker AVIII spectrometer. The typical 90° pulse lengths were 2.3 μs for ^1^H and 3.0 μs for ^31^P. The ^1^H-^31^P cross-polarization employed a linear amplitude ramp of 90-110% on ^1^H, and the center of the ramp matched to Hartmann-Hahn conditions at the first spinning sideband, with contact times of 3.5 ms and 2.5 ms for out and back CP transfers, respectively. The MAS frequency was 40 kHz controlled to within ± 10 Hz by a Bruker MAS controller. The actual sample temperature was maintained at 4 ± 1 °C throughout the experiments otherwise specified using the Bruker temperature controller.

### Solution NMR spectroscopy

1D ^1^H and ^13^C Solution NMR spectra of BVM/DMSO-D_6_ and IP6/D_2_O was collected at 14.1 T (^1^H Larmor frequency of 600.1 MHz) Bruker Avance spectrometer using a triple-resonance inverse detection (TXI) probe. ^1^H NMR spectral of BVM and IP6 are shown in Supplementary Fig. 10 and 13.

### Data processing

All MAS NMR data processed using Bruker TopSpin and NMRpipe^33^. The ^13^C and ^15^N signals were referenced with respect to the external standards adamantane and ammonium chloride respectively. ^1^H was referenced to water peak at 4.7 ppm. ^31^P was referenced with respect to the 85% H_3_PO_4_. The 2D and 3D data sets were processed by applying 30°, 45°, 60° and 90° shifted sine bell apodization followed by a Lorentzian-to-Gaussian transformation was applied in both dimensions. Forward linear prediction to twice the number of the original data points was used in the indirect dimension in some data sets followed by zero filling. The 2D CH HETCOR and dREDOR-HETCOR data of U-^13^C,^15^N,^2^H-CA_CTD_-SP1 samples were processed with gaussian window multiplication and quadrature baseline correction.

### MAS NMR chemical shift and distance restraints assignment

All the spectra were analyzed using CCPN^34^ and NMRFAM-Sparky^35,36^. The superposition of U-^13^C,^15^N-CA_CTD_-SP1/BVM/IP6, U-^13^C,^15^N-CA_CTD_-SP1/IP6 and, U-^13^C,^15^N-CA_CTD_-SP1/BVM/SO_4_ 2D CORD and 2D NCACX spectra at 50 ms shows no significant chemical shift perturbation (Supplementary figure 2). Chemical shift assignments (intra-residue/sequential assignments) were performed *de novo* on the CA_CTD_-SP1 crystalline arrays using numerous solid state NMR data sets of U-^13^C,^15^N-CA_CTD_-SP1/BVM/SO_4_: 2D CORD, 2D and 3D dipolar-based NCACX and NCOCX, 3D CONCA, and J-based 2D direct-INADEQUATE. The backbone ^15^N and carbonyl ^13^C chemical shifts assignments of residues in SP1 tail were performed on the basis of the 2D NCACX and NCOCX spectra of U-^13^C,^15^N-CA_CTD_-SP1/IP6 crystalline array at low temperature (−37 °C and -79 °C). The *de novo* assignments of inter-residue ^13^C-^13^C, ^15^N-^13^C correlations were obtained for U-,^13^C,^15^N-CA_CTD_-SP1/BVM/IP6 crystalline array using 100 ms, 250 ms, 500 ms 2D CORD spectra and 50 ms 2D NCACX and PAIN-CP. The ^13^C-^1^H and ^15^N-^1^H inter-residue correlations were obtained using ^1^H-detected CH and NH HETCOR spectra, respectively. On the basis of all spectra, for 64 residues all ^13^C and ^15^N backbone and side chain resonances were assigned and for 92 residues complete backbone assignments were obtained. For another 28 residues, complete backbone and partial side chain shift assignments were achieved and 6 more residues had partial backbone and partial side chain assignments. For 4 residues, P147, R173, V181, and M245 (SP1-M14), no resonance assignments are available since the corresponding peaks were either missing due to dynamic disorder or overlapped with other resonances. For 84 residues, amide proton assignments (HN) were completed and, of these, for 29 residues backbone proton assignments (HN and Hα) were also obtained. Additionally, multiple sidechain proton chemical shifts of various residues were assigned unambiguously.

BVM and IP6 proton correlation to proteins were assigned in 2D HC CP HETCOR, dREDOR-HETCOR, and 2D (H)PH HETCOR spectra of U-^13^C,^15^N,^2^H-CA_CTD_-SP1/BVM/IP6 and U-^13^C,^15^N,^2^H-CA_CTD_-SP1/IP6. All the samples and their experimental conditions are summarized in Supplementary Table 1. Number of cross peaks assigned in various spectra for each sample are summarized in Table 1. MAS NMR chemical shifts of CA_CTD_-SP1 crystalline array are summarized in Supplementary Table 2.

### Determination of force field parameters for BVM and IP6

The initial force field parameters of BVM and IP6 were derived by analogy following the CGENFF protocol^37^. The penalties for derived IP6 parameters and charges were all less than ten, indicating good analogy with the available atom types present in CGENFF^37^. Thus, the IP6 CGENFF parameters were directly used in structural calculations. The partial charges and bonded interactions of BVM (Supplementary Fig. 14a and 14b) with a penalty greater than ten were refined using the standard Force Field Toolkit^38^ protocol in VMD^39^ and Gaussian09^40^. Firstly, two fragments of BVM containing the high penalty scores (> 10) were generated (Supplementary Fig. 14c). In the fragments, carbon atoms at the cut points were capped with methyl groups. After fragmentation, the structures of fragments were optimized in Gaussian^40^. Then, as shown in Supplementary Fig. 14d, the partial charges in BVM were optimized to reproduce the hydrogen bond interactions with a water molecule placed nearby. Subsequently, Gaussian calculations of the hessian were used to optimize the parameters of bonds and angles^38^ with high penalty scores in the fragments. After that, torsion angle scans of the dihedral angles with large penalties were performed to generate QM target data. The dihedral angle parameters were optimized (Supplementary Fig. 14e and Supplementary Fig. 14f) towards the QM target data using the downhill Simplex method^41^. Gaussian QM energies here were calculated at the MP2/6-31G* level of theory^38^.

### Structure calculation of CA_CTD_-SP1 crystalline arrays in complex with BVM and IP6

Simulated annealing structure calculations were performed for the hexamer-of-hexamers (7 hexamer units, 42 chains) of CA_CTD_-SP1 with MAS NMR distance and dihedral restraints using Xplor-NIH version 2.53^42-44^.

#### Calculation requisites and preparation

Input protein coordinates for the calculation were obtained from X-ray crystallography (PDB ID 5I4T)^13^. The coordinates were expanded from a single hexamer-unit to a hexamer-of-hexamers containing 7 hexamer units (42 chains) using the symex command in PyMol^45^. The input coordinates of the 7 BVM and 7 IP6 molecules were from X-ray crystal structures of the molecules individually.

The distance restraints were obtained from assigned cross-peaks in MAS-NMR spectra. Both unambiguous and ambiguous restraints were considered, however, ambiguity exceeding 5-fold ambiguity were not considered. Protein-protein restraints took the form of ^13^C-^13^C, ^15^N-^13^C, ^15^N-^1^H, and ^13^C-^1^H. IP6 and BVM restraints took the form of ^1^H(IP6, BVM)-^13^C(protein) restraints and ^31^P(IP6)-^1^H(protein) restraints. The bounds of the distances are summarized in Table 2 and are consistent with our previous studies^46^. The Φ and Ψ dihedral restraints were predicted from TALOS-N^47^ with the experimental solid-state ^13^C and ^15^N chemical shifts.

For CA_CTD_-SP1-V7A/BVM/IP6 structure calculation, SP1-V7 position was mutated with alanine residue. The WT inter-residue ^13^C-^13^C, ^15^N-^13^C, and ^15^N-^1^H distance restraints along with intra-residue ^13^C-^13^C, ^13^C-^1^H, ^31^P(IP6)-^1^H(protein) distance restraints which were obtained exclusively from 2D CORD, 2D HC CP HETCOR, and 2D (H)PH HETCOR spectra of U-^13^C,^15^N,^2^H-CA_CTD_-SP1-V7A/BVM/IP6 respectively.

Prior to dynamics, the input structure underwent preparations using modules and protocols in Xplor-NIH. First, the PSF (protein structure file) was generated from a sequence file of a single chain of a hexamer unit and expanded to the 42 chains (7 hexamer units) using psfGen.duplicateSegment. The X-ray coordinates of the system were then read with protocol.initCoords and residues missing due to lacking X-ray resolution (residues 144-147, 239-245) were added as extended structures using the protocol.addUnknownAtoms routine.

Topology and parameters for both the BVM and IP6 molecules were calculated using quantum mechanical calculations, discussed in the previous section, were read in.

#### Simulation details

After reading the input coordinates and restraint files 100 structures underwent torsion angle dynamics with an annealing schedule and a final gradient minimization in Cartesian space. The BVM and IP6 molecules were free to move as rigid bodies during dynamics and final minimization.

Two identical runs of simulated annealing starting at 3000 K were performed for 10 ps, with a time step of 1 fs. The initial velocities were randomized to achieve a Maxwell distribution at a starting temperature of 3000 K. The temperature was subsequently reduced to 25 K in steps of 25 K. At each temperature step, dynamics was run for 400 fs with an initial time step of 1 fs.

#### Potential energy terms

Standard terms for bond lengths, bond angles, and improper angles were used to enforce proper covalent geometry. Standard potentials were used to incorporate distance and dihedral restraints.

A cross-correlation probability distribution potential often utilized for experimental Cryo-EM density^48^ enforced/conceded the overall shape and boundary of the crystalline array for a hexamer of hexamers. Here, an 8-Å envelope in the form of a molecular map was generated from the X-ray coordinates using the “molmap” command in UCSF Chimera 1.13^49^. To allow sidechain orientations to be defined exclusively from NMR distance and dihedral restraints the potential was restricted to the backbone atoms (N, C, CA, and O) of residues defined in the initial input X-ray coordinates.

A statistical torsion-angle potential^50^ was employed, and the gyration volume term was not included to avoid conflicts with the cross-correlation density potential. A hydrogen-bond database term, HBPot, was used to improve hydrogen-bond geometries^51^. Approximate non-crystallographic symmetry was imposed using Xplor-NIH’s PosDiffPot term, allowing the subunits of the hexamer to differ by up to 1 Å.

Force constants for distance restraints were ramped from 2 to 30 kcal/mol•Å^2^. The dihedral restraint force constants were set to 10 kcal/mol•rad^2^ for high temperature dynamics at 3000 K and 200 kcal/mol•rad^2^ during cooling. The force constants of the cross-correlation probability distribution potential were set to 50 kcal/mol during high temperature dynamics and cooling.

#### Final minimization and ensemble

After the high-temperature dynamics and cooling in dihedral space, the annealed structures were minimized using a Powell energy minimization scheme in Cartesian space. The final MAS NMR bundle comprised the 10 lowest-energy structures of the 100 calculated.

RMSD values were calculated using routines in the Xplor-NIH (version 2.51)^42-44^. The visualizations of structural elements were batch rendered in PyMOL using in-house shell/bash scripts. Secondary structure elements were classified according to TALOS-N.

## ACKNOWLEDGMENTS

This work was supported by the National Institutes of Health (NIH Grant P50-AI1504817 to A. M. G. and T. P.; R01-AI129678 to B. K. G.-P. and O. P.). We acknowledge the support of the National Science Foundation (NSF Grant CHE0959496) for the acquisition of the 850 MHz NMR spectrometer and of the National Institutes of Health (NIH Grant P30-GM110758) for the support of core instrumentation infrastructure at the University of Delaware; and of the National Institutes of Health (NIH Grant S10-OD012213) for the acquisition of the 750 MHz NMR spectrometer at the University of Pittsburgh.

## AUTHOR CONTRIBUTIONS

T. P., A. M. G., B. K. G.-P., and O. P. conceived the project and guided the work. S. S. and R. Z. performed NMR experiments and analyzed the experimental data. K. K. Z. prepared the samples. R. W. R. and S. S. performed the structure calculations. J. R. P. designed and guided the MD simulations. C. X. and J. R. P. parameterized the force fields for BVM. S. S., R. W. R., R. Z., C. X., and K. K. Z. prepared figures for the manuscript. R. W. R. wrote scripts for structure calculations, analysis of calculation results and structure visualizations. C. M. Q. took part in the design or analysis of NMR experiments. E. O. F. provided A1V and V7A viral sequence polymorphs and critical feedback on the data analysis. A. K. and S. A. performed infectivity and BVM binding studies of BVM-resistant variants. T. P., A. M. G., B. K. G.-P., and O. P. took the lead in writing the manuscript. All authors discussed the results and contributed to the manuscript preparation.

## DATA AVAILABILITY

The MAS NMR atomic structure coordinates of single hexamer of CA_CTD_-SP1/BVM/IP6 and CA_CTD_-SP1/IP6 have been deposited in the Protein Data Bank under accession code 7R7P and 7R7Q respectively. MAS NMR chemical shifts of CA_CTD_-SP1/BVM/IP6 and CA_CTD_-SP1/IP6 have been deposited in the Biological Magnetic Resonance Data Bank under accession codes 30929 and 30930.

