## Supplementary information file for "Structural Basis of HIV-1 Maturation Inhibitor Binding and Activity"

*[1]Department of Chemistry and Biochemistry, University of Delaware, Newark, DE 19716, United States; [2]Pittsburgh Center for HIV Protein Interactions, University of Pittsburgh School of Medicine, 1051 Biomedical Science Tower 3, 3501 Fifth Ave., Pittsburgh, PA 15261, United States; [3]Department of Structural Biology, University of Pittsburgh School of Medicine, 3501 Fifth Ave., Pittsburgh, PA 15261, United States; [4]HIV Dynamics and Replication Program, Center for Cancer Research, National Cancer Institute at Frederick, MD 21702-1201, United States; [5]Department of Molecular Physiology and Biological Physics, University of Virginia School of Medicine, Charlottesville, VA 22908, United States.*

***Corresponding authors:** Tatyana Polenova, Department of Chemistry and Biochemistry, University of Delaware, Newark, DE 19716, USA,; Angela M. Gronenborn, Department of Structural Biology, University of Pittsburgh School of Medicine, 3501 Fifth Ave., Pittsburgh, PA 15260, USA,; Barbie K. Ganser-Pornillos, University of Virginia School of Medicine, Charlottesville, VA 22908, USA,; Owen Pornillos, University of Virginia School of Medicine, Charlottesville, VA 22908, USA,

**Keywords:** magic angle spinning NMR, HIV-1 capsid, maturation inhibitors, Bevirimat, HIV-AIDS, atomic-resolution structure

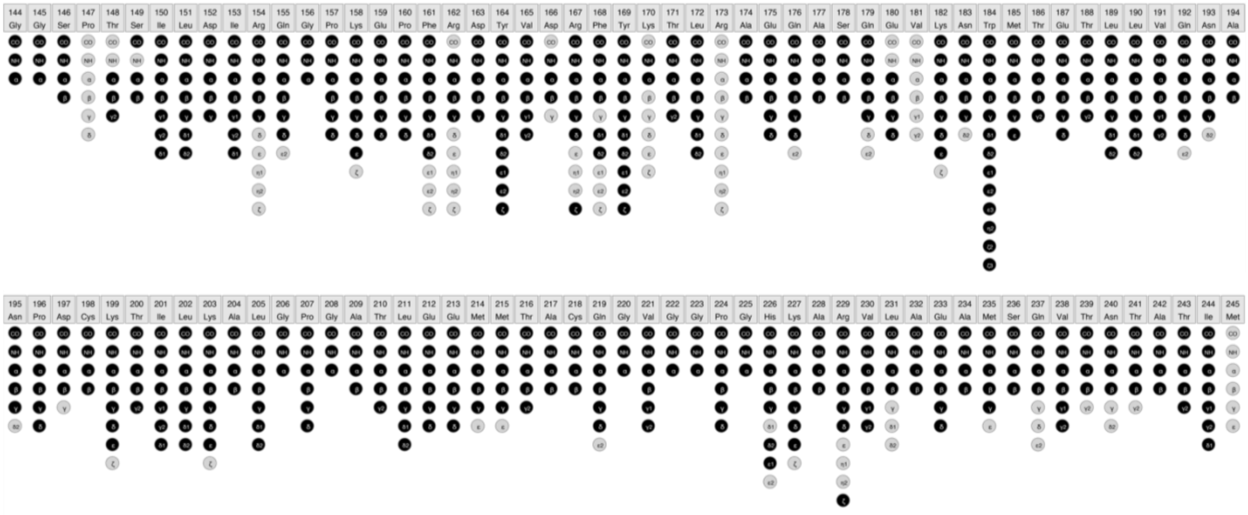

**Supplementary Figure 1: Graphical summary of resonance assignments in CA$_{CTD}$-SP1 crystalline arrays.** Atoms with assigned resonances are shown in black circles and those without assigned resonances are shown in gray circles.

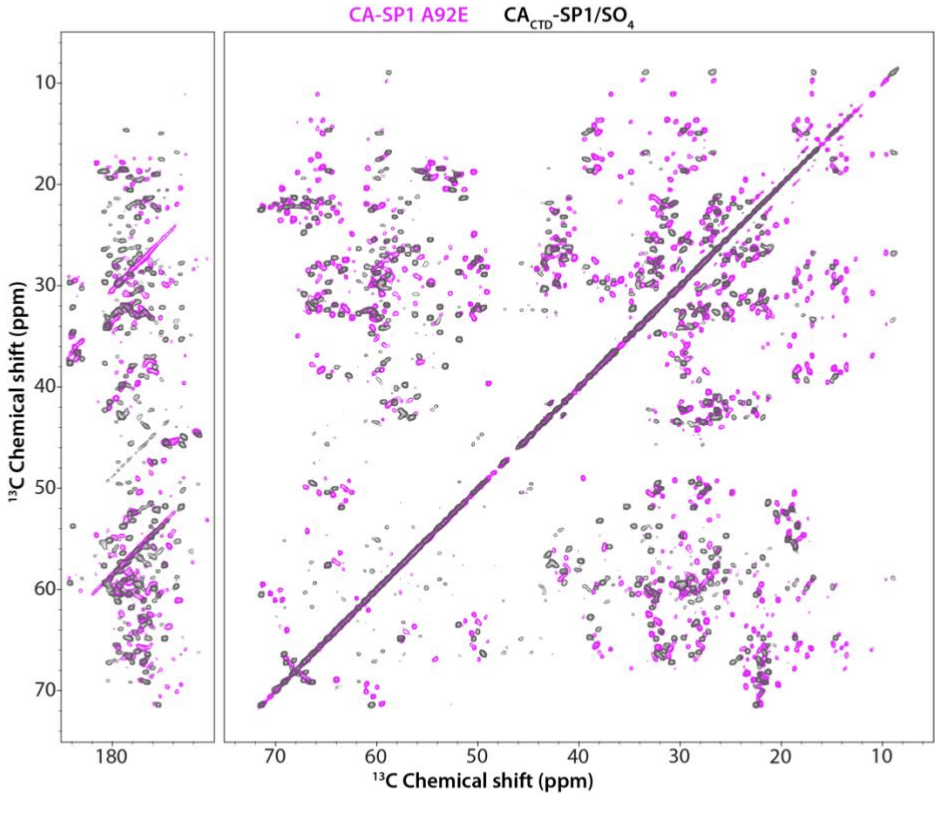


**Supplementary Figure 2: Superposition of the 2D CORD spectra of CA-SP1 tubular assemblies and CA$_{CTD}$-SP1 crystalline arrays.** The spectra were recorded at 20.0 T, with a MAS frequency of 14 kHz, and a CORD mixing time of 50 ms.

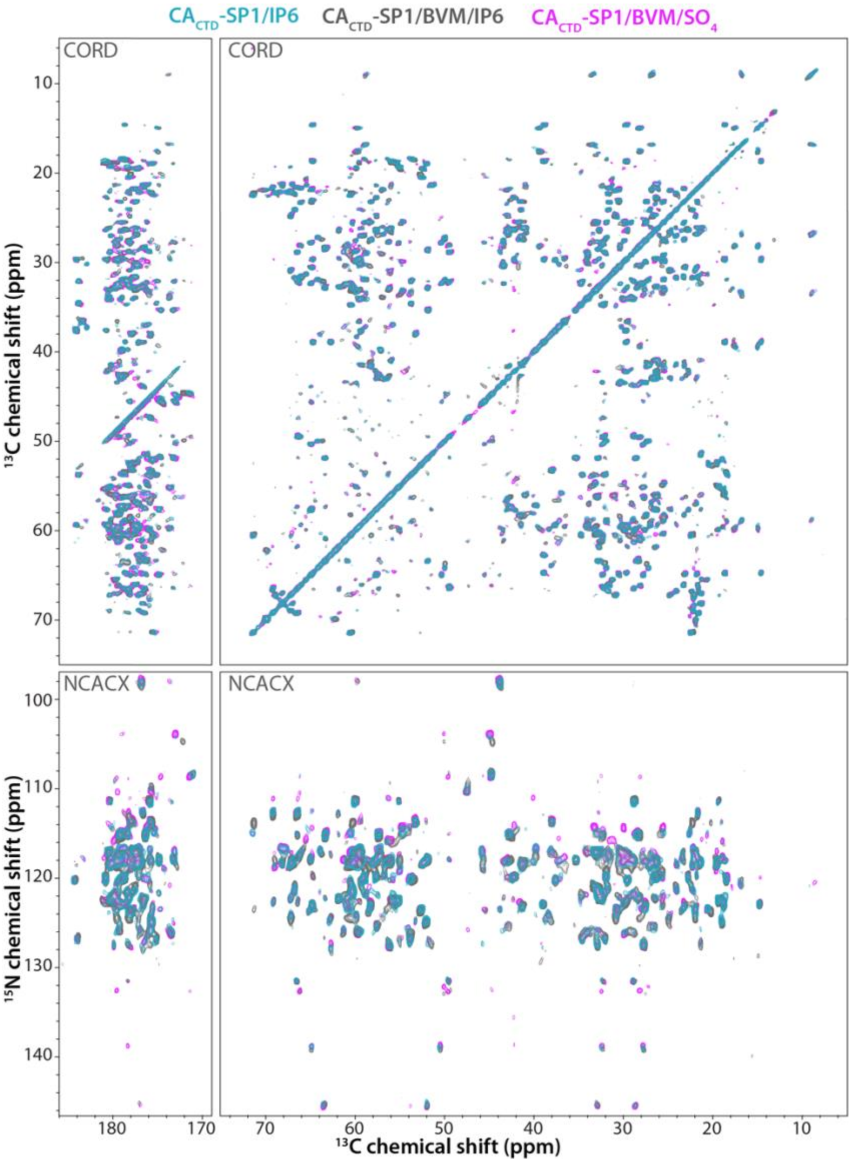


**Supplementary Figure 3: Superposition of 2D CORD (top) and 2D NCACX (bottom) spectra of U-$^{13}$C,$^{15}$N-CA$_{CTD}$-SP1/IP6, U-$^{13}$C,$^{15}$N-CA$_{CTD}$-SP1/BVM/IP6, and U-$^{13}$C,$^{15}$N-CA$_{CTD}$-SP1/BVM/SO$_4$ crystalline arrays.** The spectra were recorded at 20.0 T, with a MAS frequency of 14 kHz, and a CORD mixing time of 50 ms.

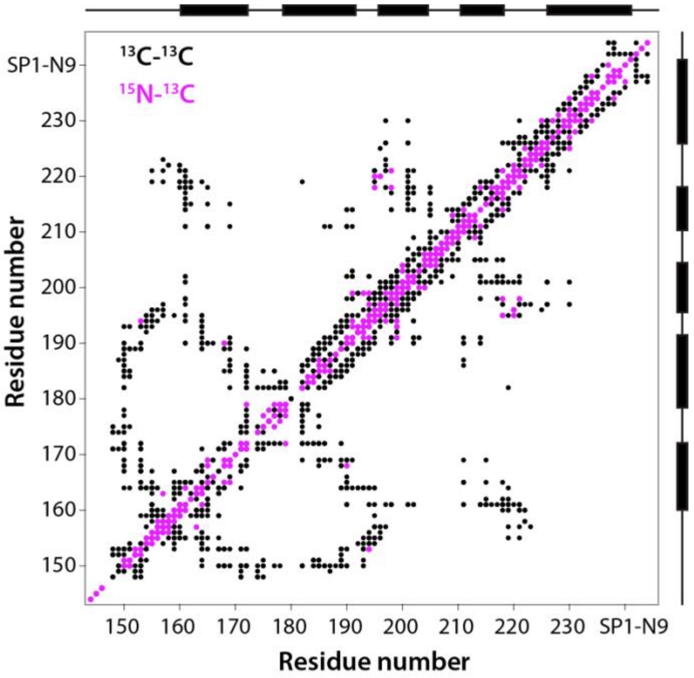


**Supplementary Figure 4: $^{15}$N-$^{13}$C and $^{13}$C-$^{13}$C contact map of BVM and IP6 bound CA$_{CTD}$-SP1 crystalline arrays.**

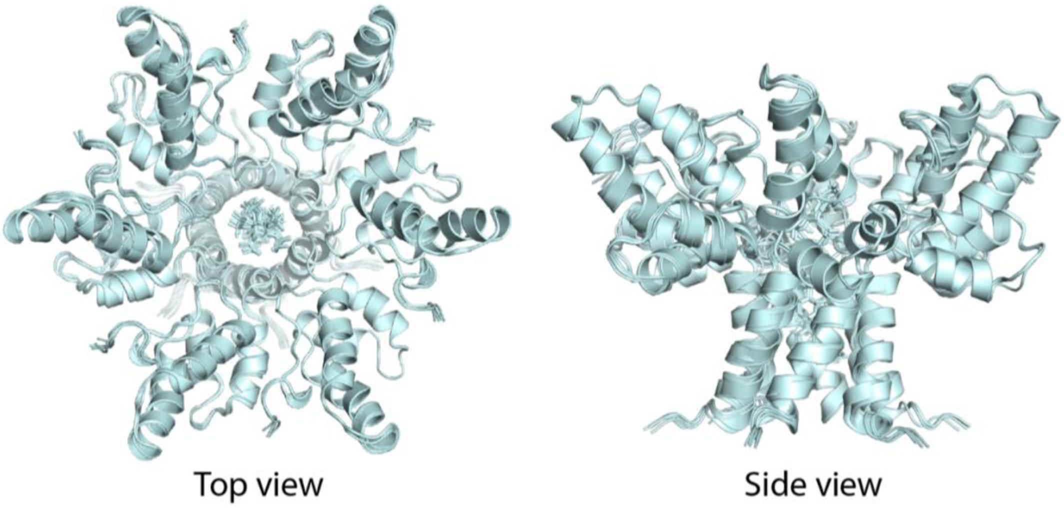


**Supplementary Figure 5: Superposition of 5 lowest energy structures of central hexamer of CA$_{CTD}$-SP1/BVM/IP6 crystalline arrays.**

| | Restraint network | $CA_{CTD}$-SP1/BVM/IP6 | $CA_{CTD}$-SP1/IP6 | | Restraint network | $CA_{CTD}$-SP1/BVM/IP6 | $CA_{CTD}$-SP1/IP6 |
|---|---|---|---|---|---|---|---|
| G144:<br>3 total restraints<br>0 long-range restraints ($\lvert i-j \rvert \geq 5$) | 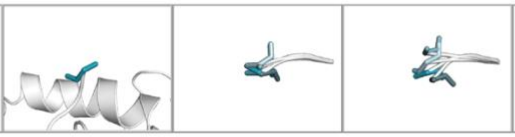 | | | G145:<br>6 total restraints<br>1 long-range restraint ($\lvert i-j \rvert \geq 5$) | 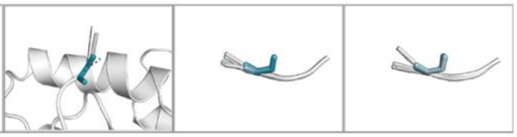 | | |
| S146:<br>8 total restraints<br>0 long-range restraints ($\lvert i-j \rvert \geq 5$) | 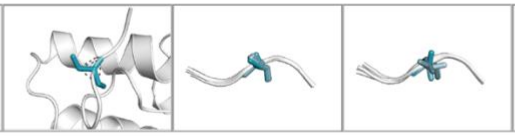 | | | P147:<br>0 total restraints<br>0 long-range restraints ($\lvert i-j \rvert \geq 5$) | 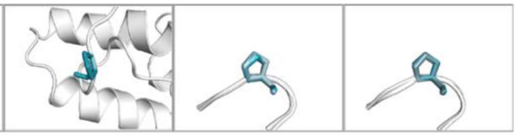 | | |
| T148:<br>20 total restraints<br>11 long-range restraints ($\lvert i-j \rvert \geq 5$) | 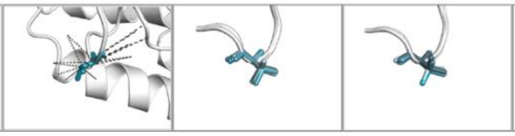 | | | S149:<br>13 total restraints<br>2 long-range restraints ($\lvert i-j \rvert \geq 5$) | 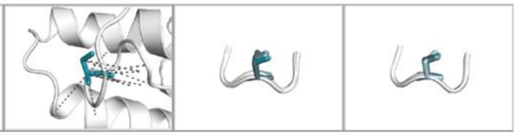 | | |
| I150:<br>112 total restraints<br>54 long-range restraints ($\lvert i-j \rvert \geq 5$) | 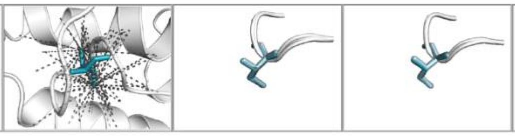 | | | L151:<br>43 total restraints<br>11 long-range restraints ($\lvert i-j \rvert \geq 5$) | 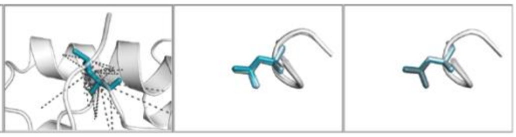 | | |
| D152:<br>28 total restraints<br>1 long-range restraint ($\lvert i-j \rvert \geq 5$) | 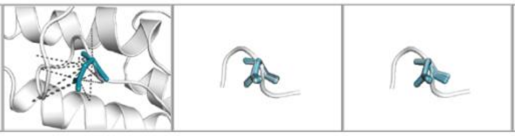 | | | I153:<br>106 total restraints<br>55 long-range restraints ($\lvert i-j \rvert \geq 5$) | 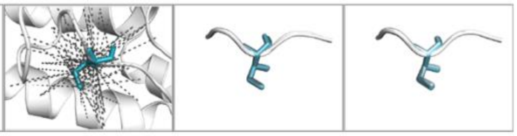 | | |
| R154:<br>33 total restraints<br>13 long-range restraints ($\lvert i-j \rvert \geq 5$) | 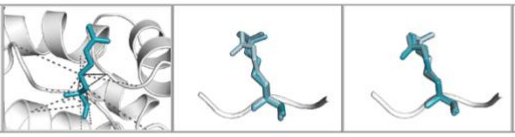 | | | Q155:<br>59 total restraints<br>31 long-range restraints ($\lvert i-j \rvert \geq 5$) | 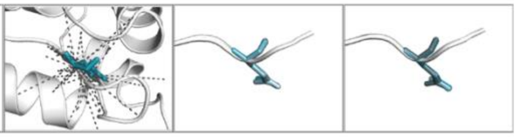 | | |
| G156:<br>32 total restraints<br>5 long-range restraints ($\lvert i-j \rvert \geq 5$) | 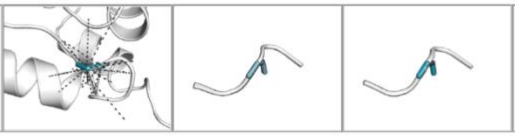 | | | P157:<br>44 total restraints<br>7 long-range restraints ($\lvert i-j \rvert \geq 5$) | 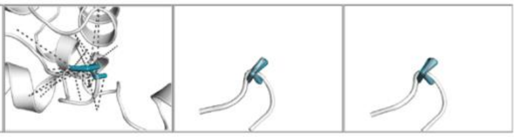 | | |
| K158:<br>45 total restraints<br>2 long-range restraints ($\lvert i-j \rvert \geq 5$) | 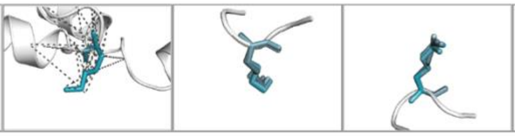 | | | E159:<br>62 total restraints<br>9 long-range restraints ($\lvert i-j \rvert \geq 5$) | 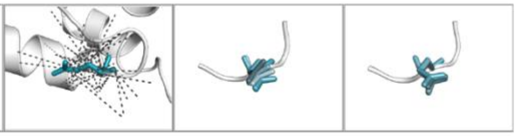 | | |
| P160:<br>55 total restraints<br>12 long-range restraints ($\lvert i-j \rvert \geq 5$) | 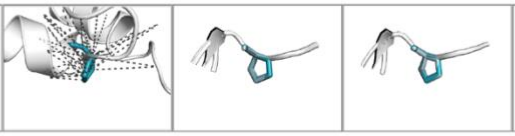 | | | F161:<br>106 total restraints<br>63 long-range restraints ($\lvert i-j \rvert \geq 5$) | 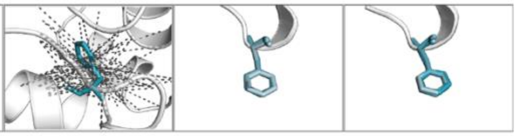 | | |
| R162:<br>12 total restraints<br>3 long-range restraints ($\lvert i-j \rvert \geq 5$) | 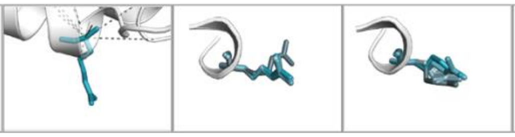 | | | D163:<br>34 total restraints<br>1 long-range restraint ($\lvert i-j \rvert \geq 5$) | 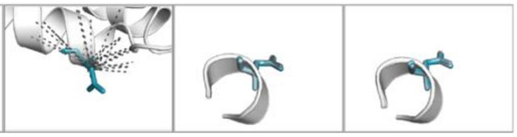 | | |
| Y164:<br>147 total restraints<br>64 long-range restraints ($\lvert i-j \rvert \geq 5$) | 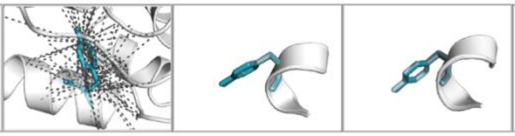 | | | V165:<br>90 total restraints<br>18 long-range restraints ($\lvert i-j \rvert \geq 5$) | 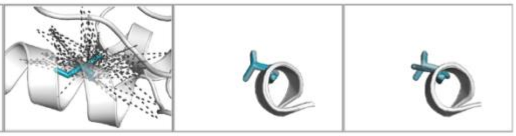 | | |
| D166:<br>8 total restraints<br>0 long-range restraints ($\lvert i-j \rvert \geq 5$) | 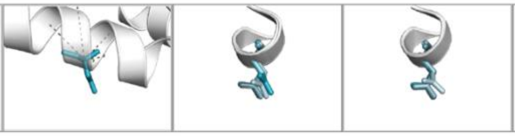 | | | R167:<br>42 total restraints<br>18 long-range restraints ($\lvert i-j \rvert \geq 5$) | 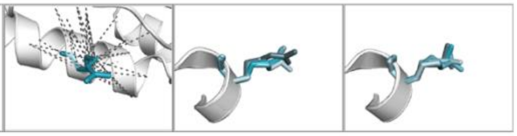 | | |

| | Restraint network | CA$_{CTD}$-SP1/BVM/IP6 | CA$_{CTD}$-SP1/IP6 | | Restraint network | CA$_{CTD}$-SP1/BVM/IP6 | CA$_{CTD}$-SP1/IP6 |
|---|---|---|---|---|---|---|---|
| F168:<br>57 total restraints<br>22 long-range restraints (\|i-j\|≥5) | 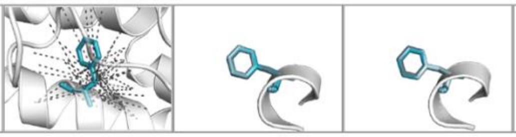 | | | Y169:<br>58 total restraints<br>15 long-range restraints (\|i-j\|≥5) | 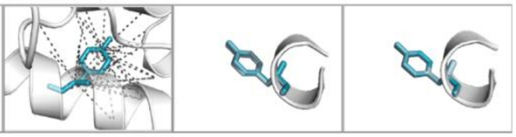 | | |
| K170:<br>1 total restraint<br>0 long-range restraints (\|i-j\|≥5) | 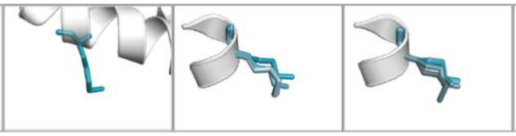 | | | T171:<br>56 total restraints<br>13 long-range restraints (\|i-j\|≥5) | 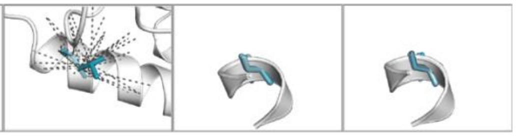 | | |
| L172:<br>68 total restraints<br>36 long-range restraints (\|i-j\|≥5) | 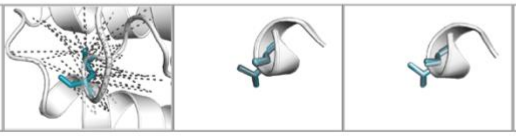 | | | R173:<br>0 total restraints<br>0 long-range restraints (\|i-j\|≥5) | 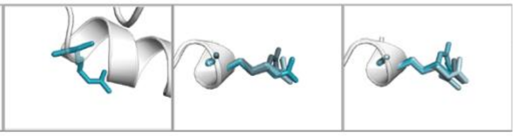 | | |
| A174:<br>23 total restraints<br>8 long-range restraints (\|i-j\|≥5) | 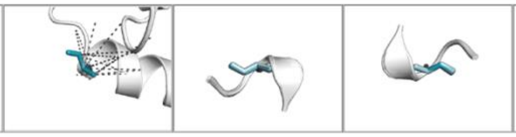 | | | E175:<br>22 total restraints<br>8 long-range restraints (\|i-j\|≥5) | 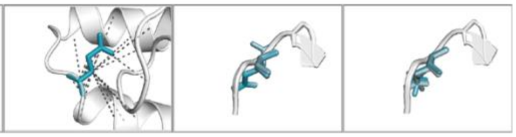 | | |
| Q176:<br>22 total restraints<br>5 long-range restraints (\|i-j\|≥5) | 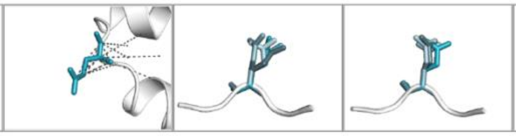 | | | A177:<br>31 total restraints<br>14 long-range restraints (\|i-j\|≥5) | 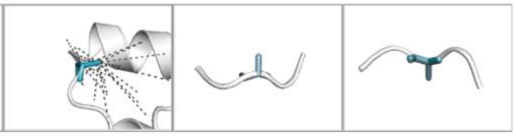 | | |
| S178:<br>25 total restraints<br>2 long-range restraints (\|i-j\|≥5) | 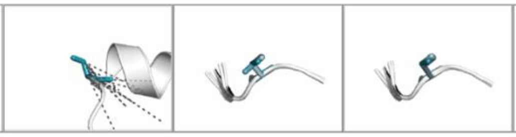 | | | Q179:<br>25 total restraints<br>0 long-range restraints (\|i-j\|≥5) | 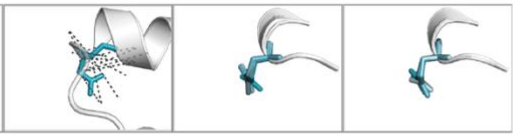 | | |
| E180:<br>6 total restraints<br>0 long-range restraints (\|i-j\|≥5) | 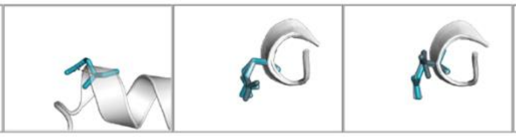 | | | V181:<br>0 total restraints<br>0 long-range restraints (\|i-j\|≥5) | 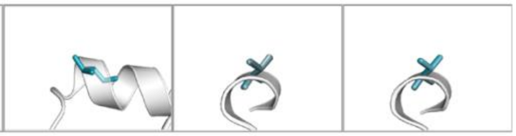 | | |
| K182:<br>69 total restraints<br>24 long-range restraints (\|i-j\|≥5) | 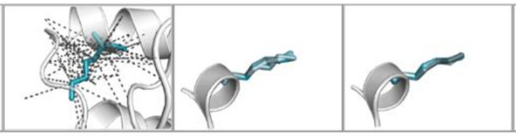 | | | N183:<br>25 total restraints<br>2 long-range restraints (\|i-j\|≥5) | 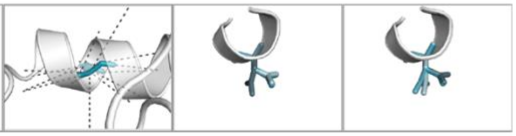 | | |
| W184:<br>61 total restraints<br>1 long-range restraint (\|i-j\|≥5) | 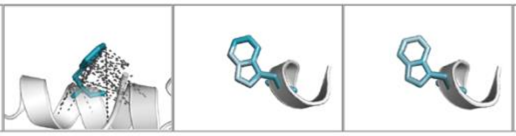 | | | M185:<br>58 total restraints<br>24 long-range restraints (\|i-j\|≥5) | 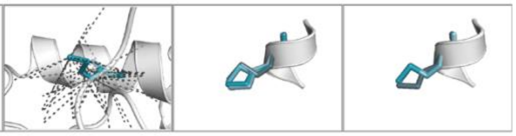 | | |
| T186:<br>41 total restraints<br>13 long-range restraints (\|i-j\|≥5) | 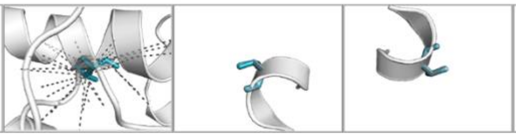 | | | E187:<br>40 total restraints<br>2 long-range restraints (\|i-j\|≥5) | 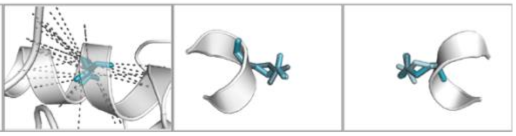 | | |
| T188:<br>34 total restraints<br>1 long-range restraint (\|i-j\|≥5) | 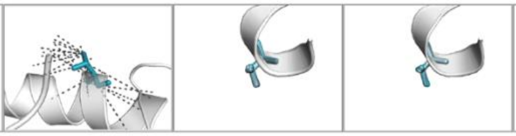 | | | L189:<br>78 total restraints<br>35 long-range restraints (\|i-j\|≥5) | 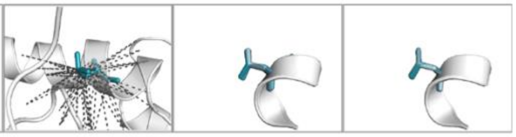 | | |
| L190:<br>96 total restraints<br>52 long-range restraints (\|i-j\|≥5) | 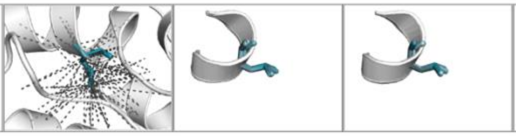 | | | V191:<br>72 total restraints<br>33 long-range restraints (\|i-j\|≥5) | 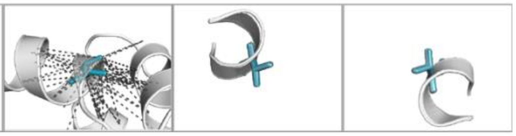 | | |

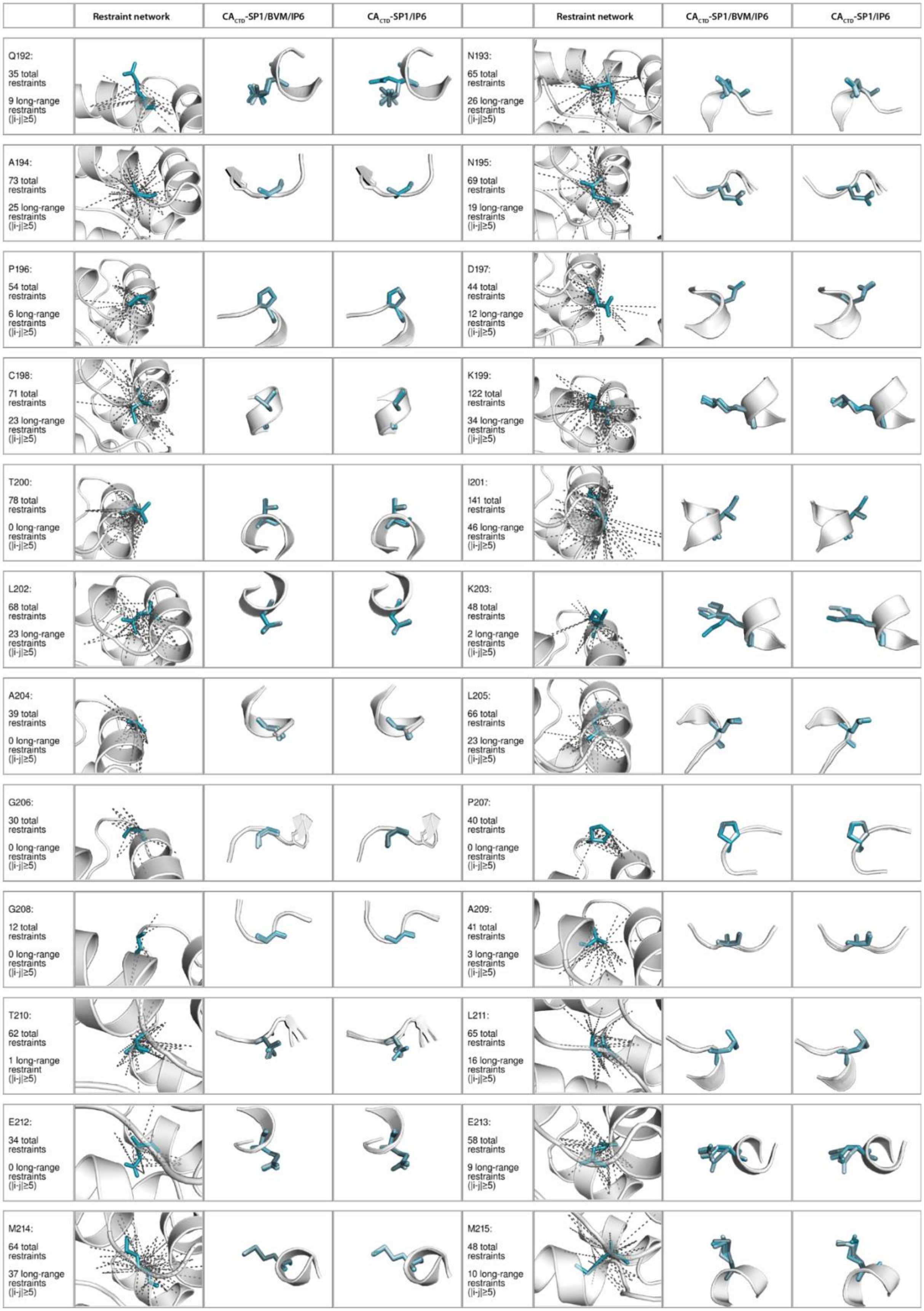
Restraint network
$CA_{CTD}$-SP1/BVM/IP6
$CA_{CTD}$-SP1/IP6
Restraint network
$CA_{CTD}$-SP1/BVM/IP6
$CA_{CTD}$-SP1/IP6
Q192: 35 total restraints 9 long-range restraints (|i-j|≥5)
N193: 65 total restraints 26 long-range restraints (|i-j|≥5)
A194: 73 total restraints 25 long-range restraints (|i-j|≥5)
N195: 69 total restraints 19 long-range restraints (|i-j|≥5)
P196: 54 total restraints 6 long-range restraints (|i-j|≥5)
D197: 44 total restraints 12 long-range restraints (|i-j|≥5)
C198: 71 total restraints 23 long-range restraints (|i-j|≥5)
K199: 122 total restraints 34 long-range restraints (|i-j|≥5)
T200: 78 total restraints 0 long-range restraints (|i-j|≥5)
I201: 141 total restraints 46 long-range restraints (|i-j|≥5)
L202: 68 total restraints 23 long-range restraints (|i-j|≥5)
K203: 48 total restraints 2 long-range restraints (|i-j|≥5)
A204: 39 total restraints 0 long-range restraints (|i-j|≥5)
L205: 66 total restraints 23 long-range restraints (|i-j|≥5)
G206: 30 total restraints 0 long-range restraints (|i-j|≥5)
P207: 40 total restraints 0 long-range restraints (|i-j|≥5)
G208: 12 total restraints 0 long-range restraints (|i-j|≥5)
A209: 41 total restraints 3 long-range restraints (|i-j|≥5)
T210: 62 total restraints 1 long-range restraint (|i-j|≥5)
L211: 65 total restraints 16 long-range restraints (|i-j|≥5)
E212: 34 total restraints 0 long-range restraints (|i-j|≥5)
E213: 58 total restraints 9 long-range restraints (|i-j|≥5)
M214: 64 total restraints 37 long-range restraints (|i-j|≥5)
M215: 48 total restraints 10 long-range restraints (|i-j|≥5)

| | Restraint network | CA$_{CTD}$-SP1/BVM/IP6 | CA$_{CTD}$-SP1/IP6 | | Restraint network | CA$_{CTD}$-SP1/BVM/IP6 | CA$_{CTD}$-SP1/IP6 |
|---|---|---|---|---|---|---|---|
| T216:<br>47 total restraints<br>2 long-range restraints (\|i-j\|≥5) | 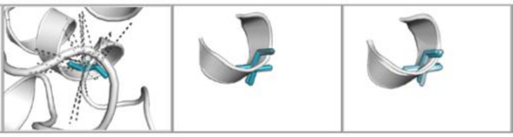 | | | A217:<br>49 total restraints<br>11 long-range restraints (\|i-j\|≥5) | 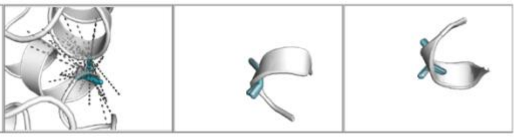 | | |
| C218:<br>73 total restraints<br>37 long-range restraints (\|i-j\|≥5) | 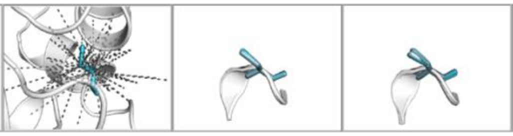 | | | Q219:<br>51 total restraints<br>12 long-range restraints (\|i-j\|≥5) | 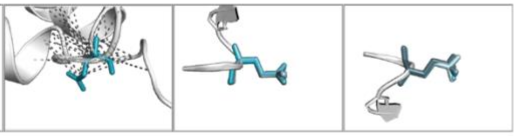 | | |
| G220:<br>33 total restraints<br>7 long-range restraints (\|i-j\|≥5) | 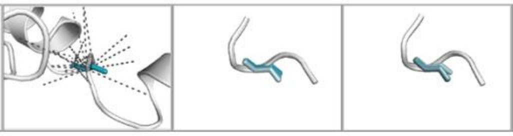 | | | V221:<br>94 total restraints<br>53 long-range restraints (\|i-j\|≥5) | 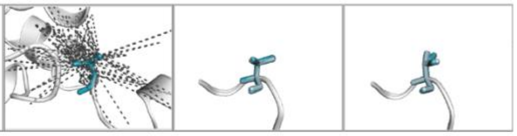 | | |
| G222:<br>23 total restraints<br>8 long-range restraints (\|i-j\|≥5) | 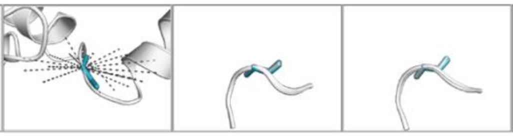 | | | G223:<br>30 total restraints<br>0 long-range restraints (\|i-j\|≥5) | 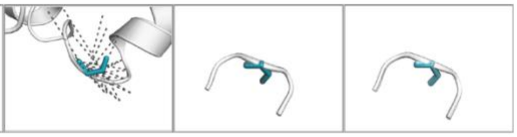 | | |
| P224:<br>55 total restraints<br>0 long-range restraints (\|i-j\|≥5) | 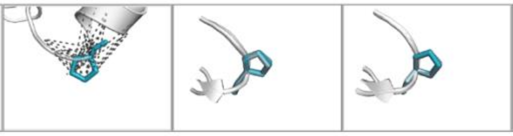 | | | G225:<br>35 total restraints<br>1 long-range restraint (\|i-j\|≥5) | 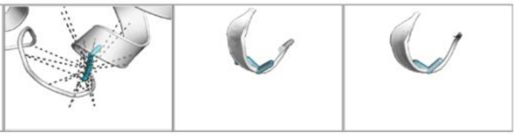 | | |
| H226:<br>109 total restraints<br>31 long-range restraints (\|i-j\|≥5) | 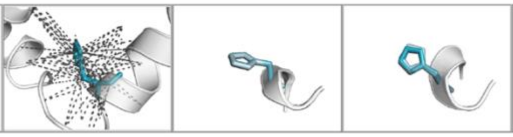 | | | K227:<br>50 total restraints<br>7 long-range restraints (\|i-j\|≥5) | 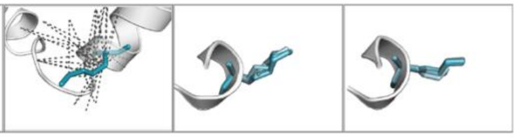 | | |
| A228:<br>28 total restraints<br>0 long-range restraints (\|i-j\|≥5) | 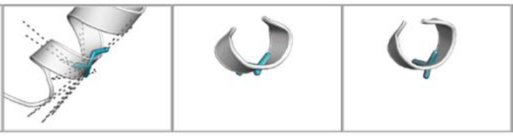 | | | R229:<br>47 total restraints<br>0 long-range restraints (\|i-j\|≥5) | 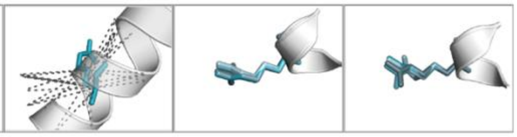 | | |
| V230:<br>63 total restraints<br>5 long-range restraints (\|i-j\|≥5) | 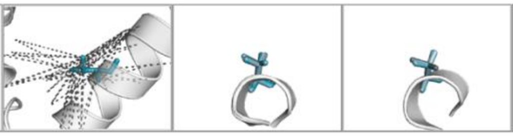 | | | L231:<br>13 total restraints<br>0 long-range restraints (\|i-j\|≥5) | 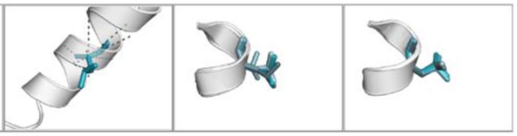 | | |
| A232:(SP1-A1)<br>31 total restraints<br>0 long-range restraints (\|i-j\|≥5) | 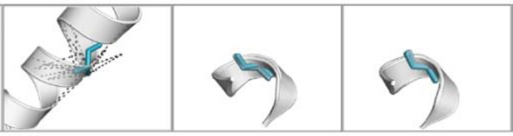 | | | E233: (SP1-E2)<br>57 total restraints<br>0 long-range restraints (\|i-j\|≥5) | 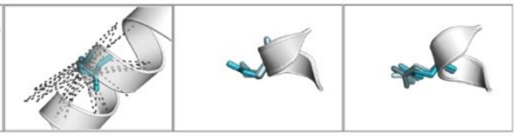 | | |
| A234:(SP1-A3)<br>36 total restraints<br>0 long-range restraints (\|i-j\|≥5) | 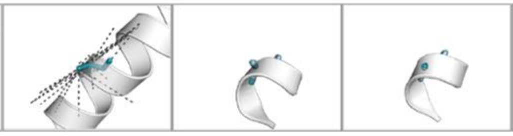 | | | M235:(SP1-M4)<br>28 total restraints<br>1 long-range restraint (\|i-j\|≥5) | 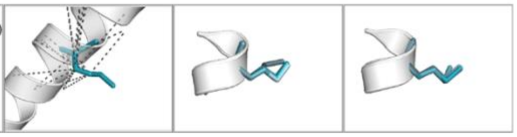 | | |
| S236:(SP1-S5)<br>11 total restraints<br>0 long-range restraints (\|i-j\|≥5) | 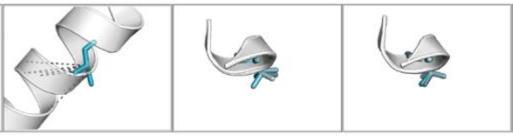 | | | Q237: (SP1-Q6)<br>24 total restraints<br>3 long-range restraints (\|i-j\|≥5) | 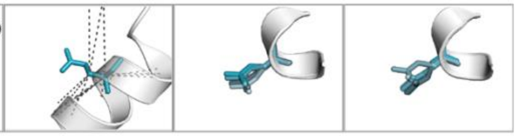 | | |
| V238:(SP1-V7)<br>38 total restraints<br>4 long-range restraints (\|i-j\|≥5) | 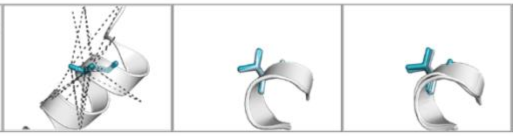 | | | T239: (SP1-T8)<br>11 total restraints<br>0 long-range restraints (\|i-j\|≥5) | 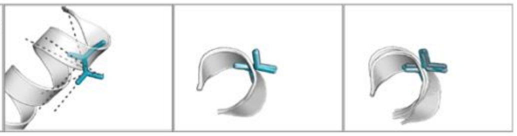 | | |

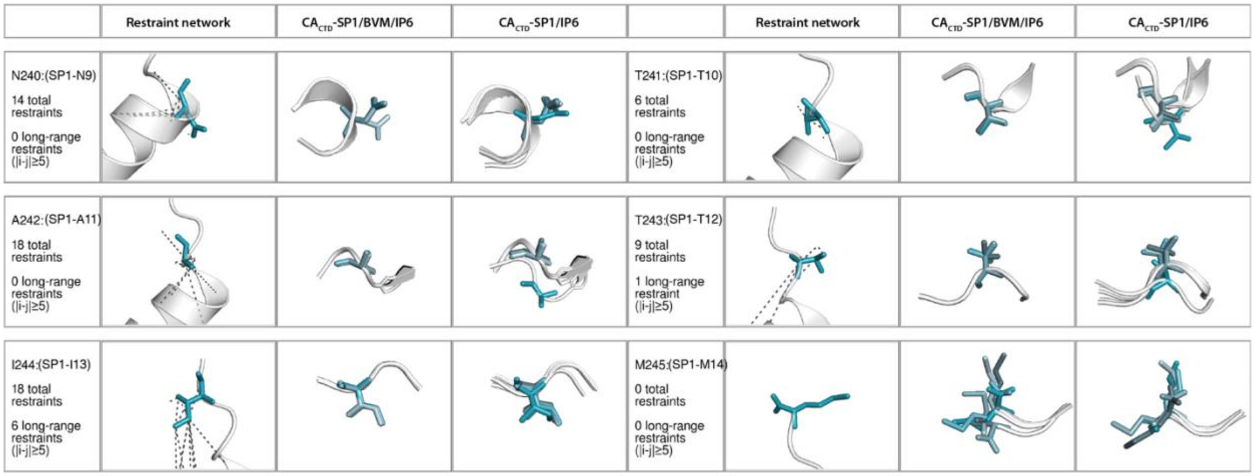


**Supplementary Figure 6: Side chain conformations in the refined structure of $CA_{CTD}$-SP1/BVM/IP6 crystalline arrays.** Experimental NMR intra-chain distances (gray dotted lines) are mapped onto the structure for each residue, if available.

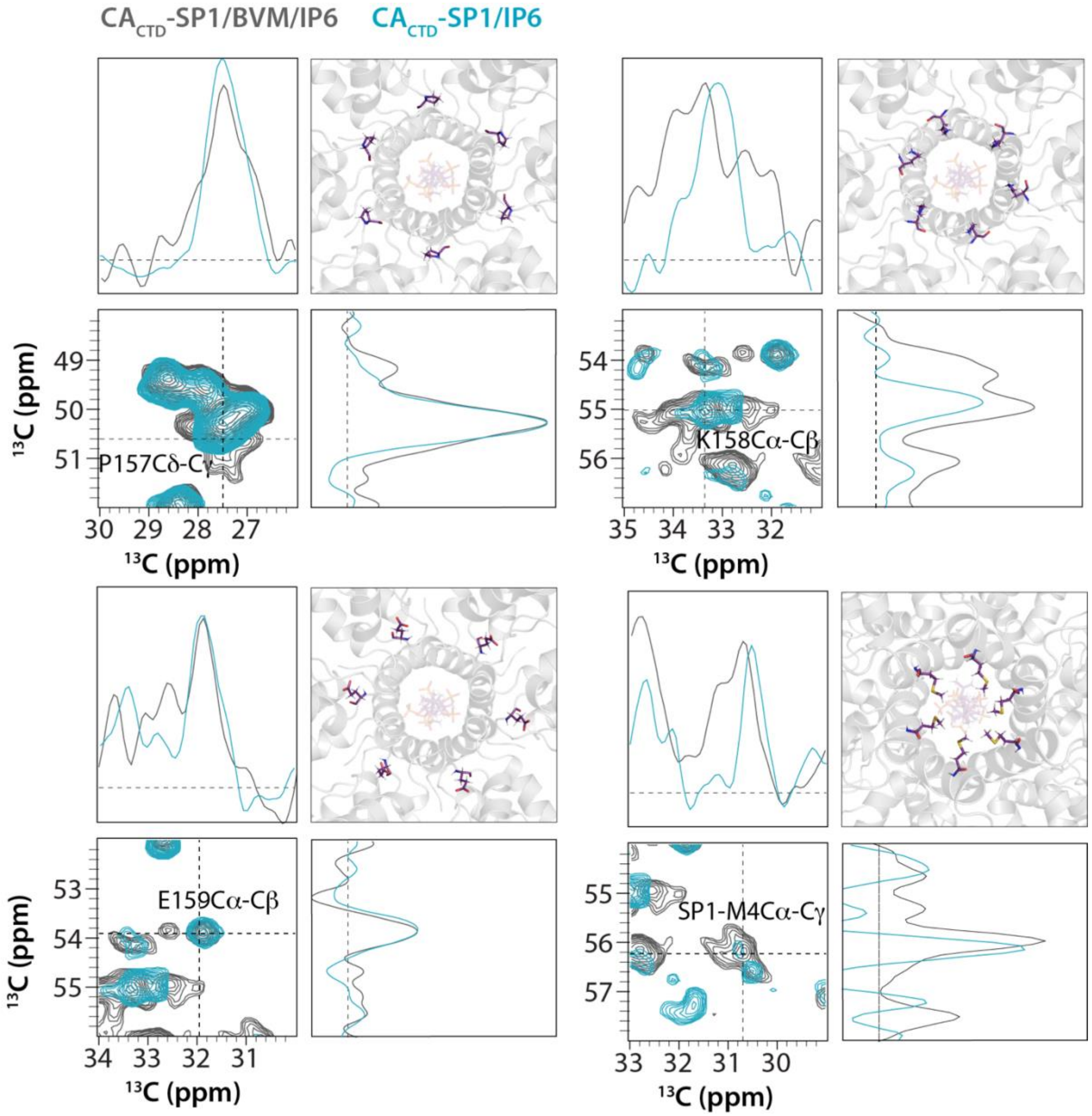


**Supplementary Figure 7: BVM binding to CA$_{CTD}$-SP1 crystalline arrays induces conformational heterogeneity in the binding region.** Expansions around selected regions of the CORD spectra of CA$_{CTD}$-SP1/IP6 (cyan) and CA$_{CTD}$-SP1/BVM/IP6 (grey). The spectra were recorded at 20.0 T, with a MAS frequency of 14 kHz, and a CORD mixing time of 50 ms.

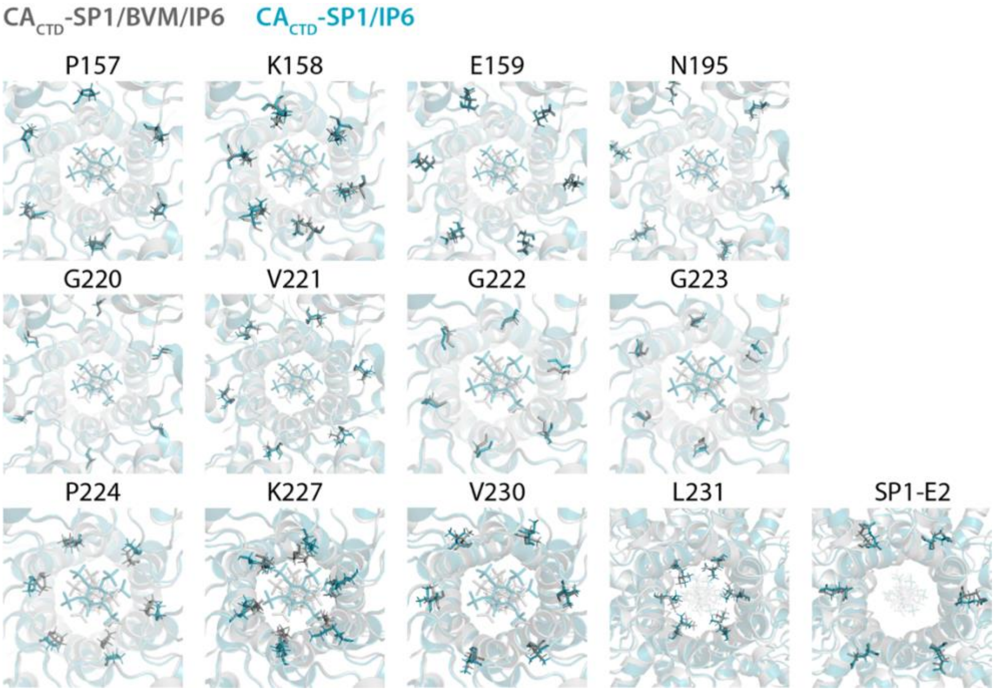


**Supplementary Figure 8: Reorientation of side chains in CA$_{CTD}$-SP1 crystalline arrays induced by BVM binding.** The sidechain conformations were determined based on multiple NMR distance restraints as described in Methods and discussed in the text.

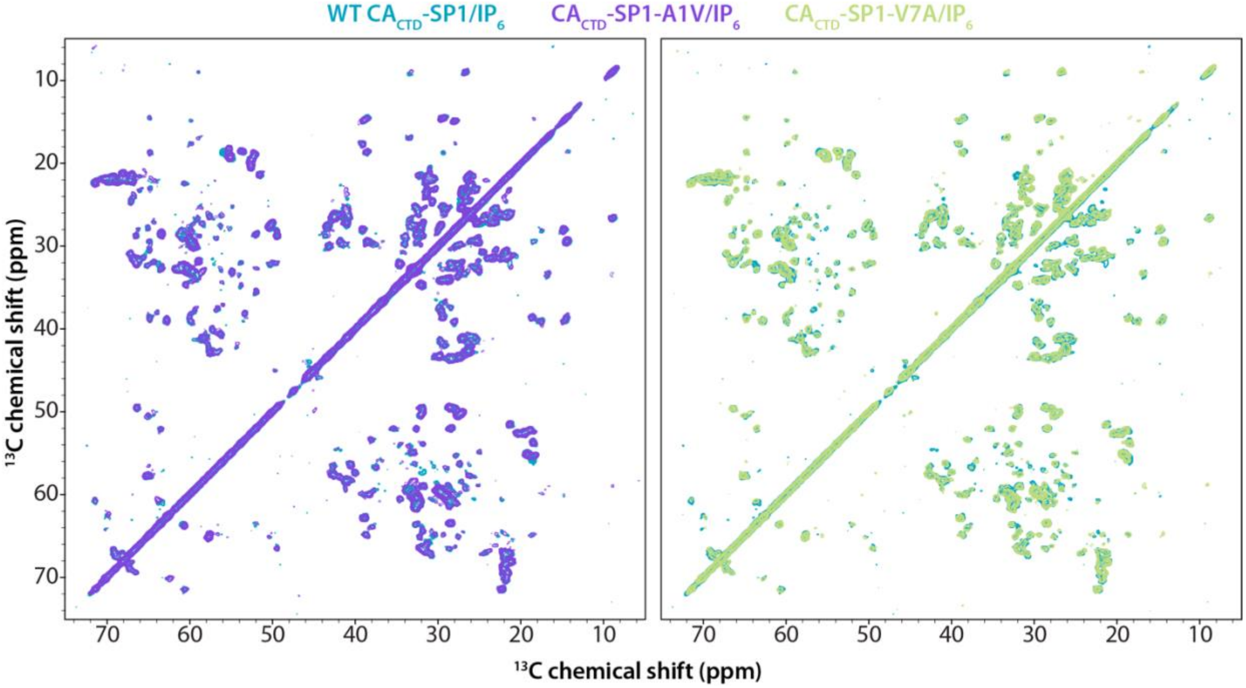


**Supplementary Figure 9: Comparison of 2D CORD spectra of wild-type, A1V, V7A variant.** Left: Superposition of 2D CORD spectra of U-$^{13}$C,$^{15}$N,$^{2}$H-CA$_{CTD}$-SP1/IP6 (cyan) and U-$^{13}$C,$^{15}$N,$^{2}$H-CA$_{CTD}$-SP1-A1V/IP6 (purple). Right: Superposition of 2D CORD spectra of U-$^{13}$C,$^{15}$N,$^{2}$H-CA$_{CTD}$-SP1/IP6 (cyan) and U-$^{13}$C,$^{15}$N,$^{2}$H-CA$_{CTD}$-SP1-V7A/IP6 (light green).

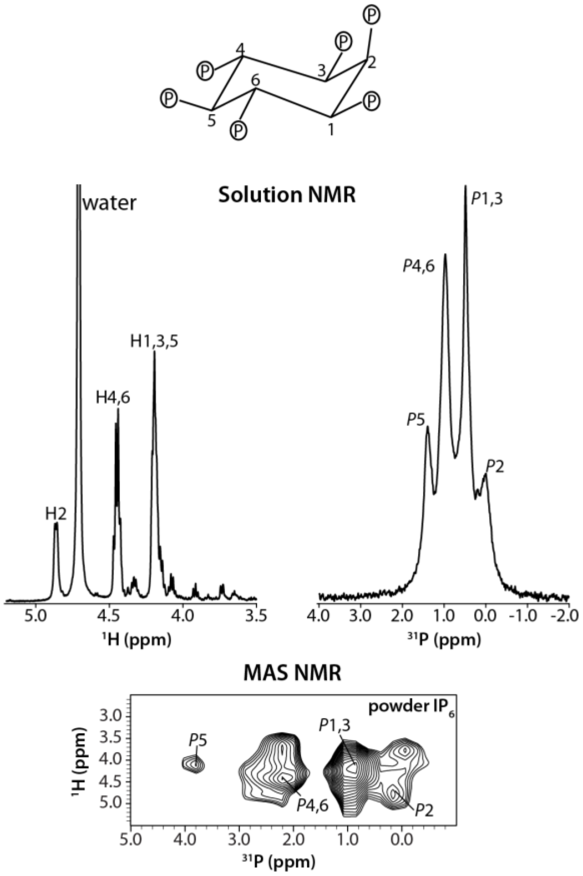


**Supplementary Figure 10: 1D solution NMR and 2D MAS NMR spectra of IP6.** 1D solution NMR spectra of IP6/$D_2O$ were recorded at 14.1 T. 2D (H)PH HETCOR spectra of powder IP6 was recorded at 20.1 T.

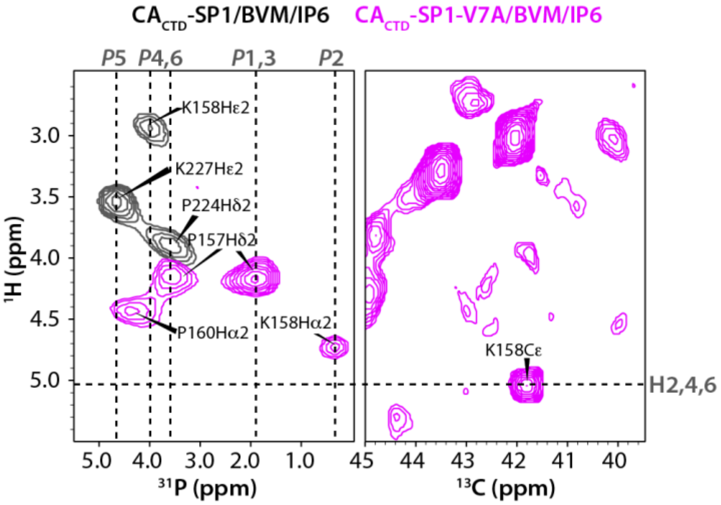


**Supplementary Figure 11: MAS NMR chemical shift assignment of IP6 bound to wild-type and V7A $CA_{CTD}$-SP1 with BVM. a)** IP6 phosphorus (*P*) and protons are labeled outside the box. The left panel is overlay of 2D (H)PH HETCOR spectra of U-$^{13}$C,$^{15}$N,$^{2}$H-$CA_{CTD}$-SP1/BVM/IP6 (grey) and U-$^{13}$C,$^{15}$N,$^{2}$H-$CA_{CTD}$-SP1-V7A/BVM/IP6 (magenta). The right panel is 2D HC HETCOR spectra of U-$^{13}$C,$^{15}$N,$^{2}$H-$CA_{CTD}$-SP1-V7A/BVM/IP6 showing the H2,4,6 (IP6) – K158Cε ($CA_{CTD}$-SP1) correlation. This indicates that in 2D (H)PH HETCOR spectra the correlations are not from intramolecular IP6 correlations.

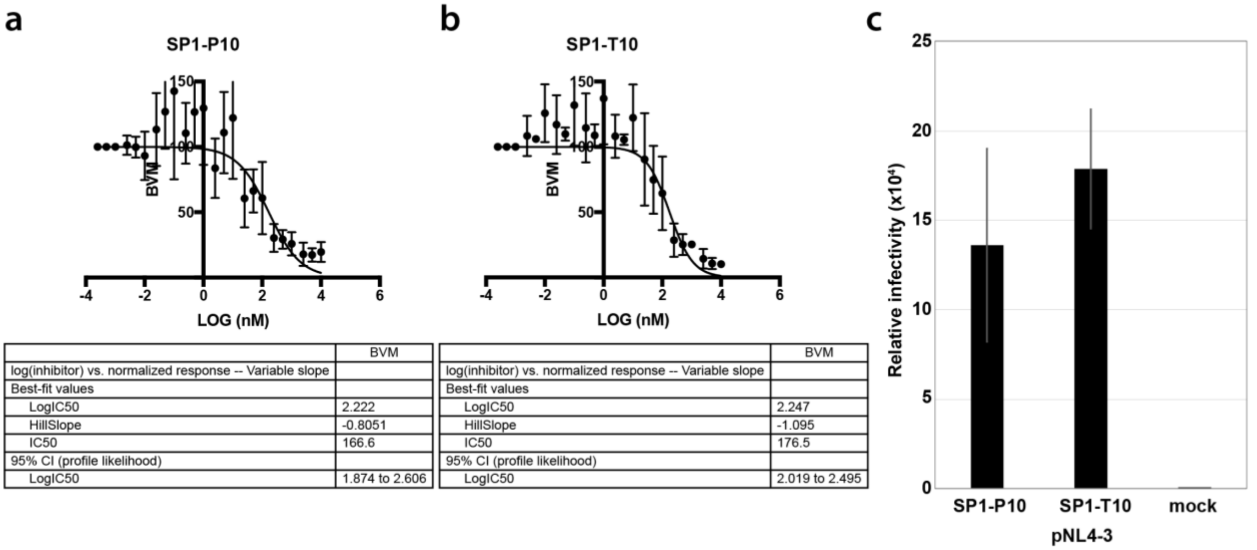


**Supplementary Figure 12: Viral infectivity and sensitivity to BVM.** Antiviral activity of BVM against **a)** SP1-P10 and **b)** the SP1-T10 sequence variants. **c)** Relative infectivity of SP1-P10 and SP1-T10 sequence variants using the TZM-bl indicator cell line.

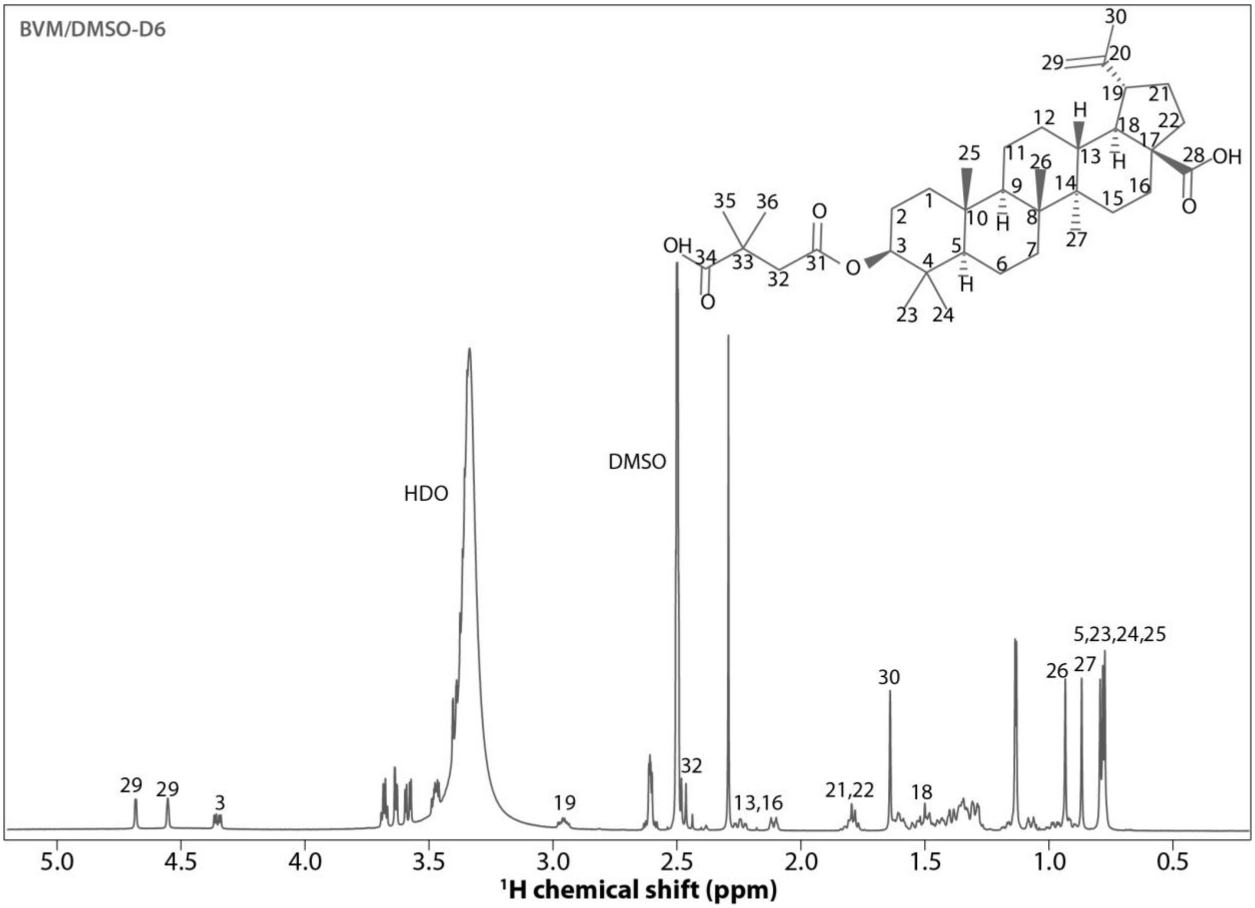


**Supplementary Figure 13: $^{1}H$ solution NMR spectra of BVM dissolved in DMSO-$D_6$ (14.1 T) .**

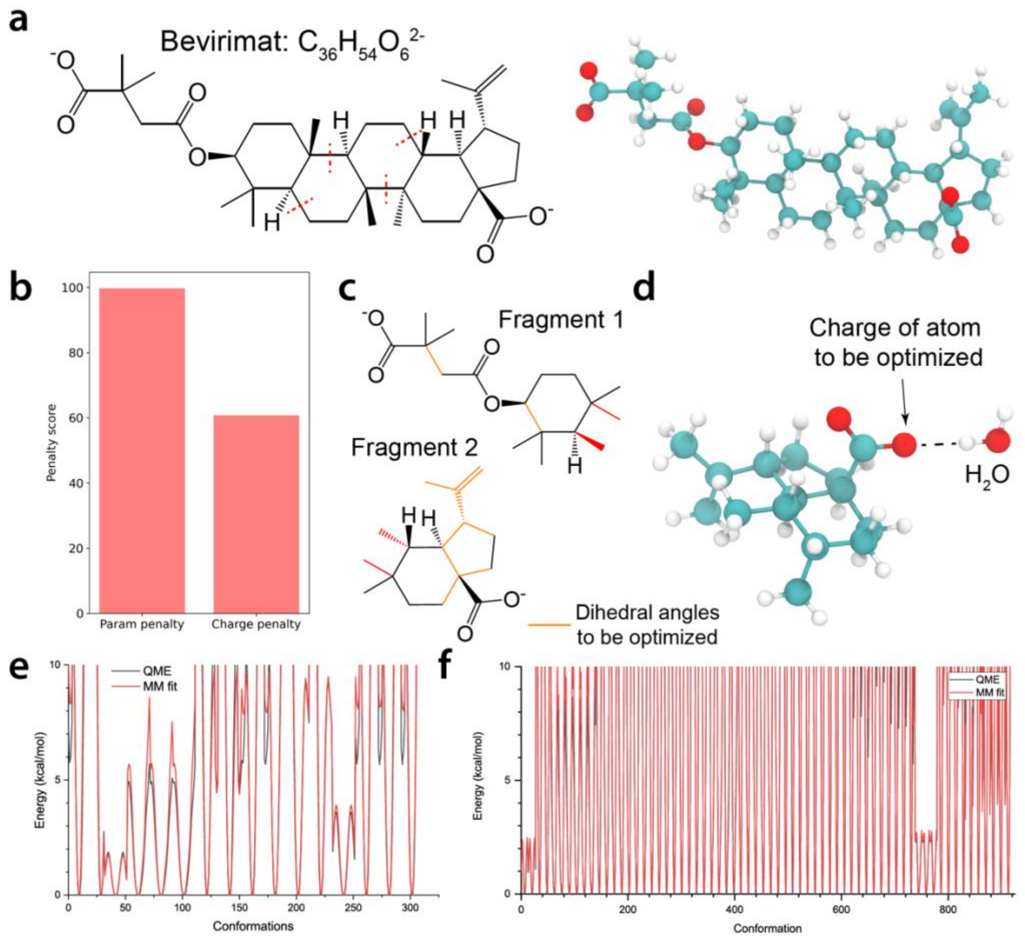


**Supplementary Figure 14: Determination of force field parameters for Bevirimat (BVM). a)** Chemical and atomic structure of BVM. **b)** Penalty scores of CGENFF generated BVM parameters. **c)** Fragmentation of BVM molecule. Two fragments of BVM contain parameters of high penalty scores were generated. Bonds colored in orange are the internal bonds of dihedral angles with a parameter penalty score larger than ten. Bonds in red represent the bonds connect to methyl groups that were used to cap the carbon at the cut points. **d)** Optimization of partial charge of atoms. A water molecule was placed close to the atom whose charge is to be optimized. **e)** and **f)** Potential energy surface profiles of the dihedral angle scans and fitting results for BVM fragment 1 and 2, respectively.

**Supplementary Table 1:** Summary of NMR experiments

| Solid State MAS NMR | | | | | | |
|---|---|---|---|---|---|---|
| **ID** | **Sample** | **Experiment** | **$B_0$ (T)** | **$\omega_r$ (kHz)** | **CC mixing time (ms)** | **T (± 1°C)** |
| 1 | U-$^{13}$C,$^{15}$N-CA$_{CTD}$-SP1/BVM/IP6 | 2D CORD | 20.0 | 14 | 10 | 4 |
| | | 2D CORD | 20.0 | 14 | 25 | 4 |
| | | 2D CORD | 20.0 | 14 | 50 | 4 |
| | | 2D CORD | 20.0 | 14 | 100 | 4 |
| | | 2D CORD | 20.0 | 14 | 200 | 4 |
| | | 2D CORD | 20.0 | 14 | 10 | -10 |
| | | 2D CORD | 20.0 | 14 | 25 | -10 |
| | | 2D CORD | 20.0 | 14 | 100 | -10 |
| | | 2D CORD | 20.0 | 14 | 200 | -10 |
| | | 2D NCACX | 20.0 | 14 | 50 | 4 |
| | | 2D NCACX | 20.0 | 14 | 50 | -10 |
| | | 2D CORD | 14.1 | 14 | 25 | 4 |
| | | 2D CORD | 14.1 | 14 | 100 | 4 |
| | | 2D CORD | 14.1 | 14 | 250 | 4 |
| | | 2D CORD | 14.1 | 14 | 500 | 4 |
| | | 2D INADEQUATE | 14.1 | 14 | | 4 |
| | | 2D PAIN CP | 14.1 | 14 | | 4 |
| 2 | U-$^{13}$C,$^{15}$N-CA$_{CTD}$-SP1/IP6 | 2D CORD | 20.1 | 14 | 50 | 4 |
| | | 2D CORD | 20.1 | 14 | 100 | 4 |
| | | 2D NCACX | 20.1 | 14 | 50 | 4 |
| | | 2D NCACX | 20.1 | 14 | 50 | -10 |
| | | 3D NCOCX | 20.1 | 14 | 25 | 4 |
| | | 2D CORD | 14.1 | 14 | 25 | 4 |
| | | 2D CORD | 14.1 | 14 | 50 | 4 |
| | | 2D NCACX | 14.1 | 14 | 25 | 4 |
| | | 2D NCOCX | 14.1 | 14 | 25 | 4 |
| | | 3D NCACX | 14.1 | 14 | 25 | 4 |
| | | 2D NCACX | 17.6 | 15 | 25 | -79 |
| | | 2D NCACX | 17.6 | 15 | 25 | -37 |
| | | 2D NCOCX | 17.6 | 15 | 25 | -79 |
| 3 | U-$^{13}$C,$^{15}$N-CA$_{CTD}$-SP1/BVM/$SO_4$ | 2D CORD | 20.1 | 14 | 10 | 4 |
| | | 2D CORD | 20.1 | 14 | 50 | 4 |
| | | 2D CORD | 20.1 | 14 | 100 | 4 |
| | | 2D CORD | 20.1 | 14 | 200 | 4 |
| | | 2D CORD | 20.1 | 14 | 10 | -10 |
| | | 2D CORD | 20.1 | 14 | 100 | -10 |
| | | 2D NCACX | 20.1 | 14 | 50 | 4 |
| | | 2D NCACX | 20.1 | 14 | 50 | -10 |

| | | | | | | |
|---|---|---|---|---|---|---|
| | | 2D INADEQUATE | 20.1 | 14 | | 4 |
| | | 2D CORD | 14.1 | 14 | 10 | 4 |
| | | 2D CORD | 14.1 | 14 | 25 | 4 |
| | | 2D NCACX | 14.1 | 14 | 25 | 4 |
| | | 2D NCACX | 14.1 | 14 | 50 | 4 |
| | | 2D NCOCX | 14.1 | 14 | 50 | 4 |
| | | 3D NCACX | 14.1 | 14 | 25 | 4 |
| | | 3D NCOCX | 14.1 | 14 | 25 | 4 |
| | | 3D CONCA | 14.1 | 14 | | 4 |
| 4 | U-$^{13}$C,$^{15}$N-CA$_{CTD}$-SP1/SO$_4$ | 2D CORD | 20.1 | 14 | 50 | 4 |
| | | 2D NCA | 20.1 | 14 | | 4 |
| | | 2D NCACX | 20.1 | 14 | 50 | 4 |
| | | 2D CORD | 14.1 | 14 | 25 | 4 |
| | | 2D NCACX | 14.1 | 14 | 25 | 4 |
| | | 2D NCOCX | 14.1 | 14 | 25 | 4 |
| | | 2D NCOCX | 14.1 | 14 | 50 | 4 |
| | | 3D NCACX | 14.1 | 14 | 25 | 4 |
| | | 2D INADEQUATE | 14.1 | 14 | | 4 |
| 5 | U-$^{13}$C,$^{15}$N,$^{2}$H-CA$_{CTD}$-SP1/BVM/IP6 | 1D dREDOR | 20.1 | 40 | | 4 |
| | | 1D $^{31}$P Direct | 20.1 | 40 | | 4 |
| | | 1D $^{31}$P CP | 20.1 | 40 | | 4 |
| | | 2D CORD | 20.1 | 14 | | -5 |
| | | 2D HC CP HETCOR | 20.1 | 40 | | 4 |
| | | 2D dREDOR-HETCOR | 20.1 | 40 | | 4 |
| | | 2D NH HETCOR | 20.1 | 40 | | 4 |
| | | 2D (H)PH HETCOR | 20.1 | 40 | | 4 |
| 6 | U-$^{13}$C,$^{15}$N,$^{2}$H-CA$_{CTD}$-SP1/IP6 (Buffer A) | 1D dREDOR | 20.1 | 40 | | 4 |
| | | 1D $^{31}$P Direct | 20.1 | 40 | | 4 |
| | | 1D $^{31}$P CP | 20.1 | 40 | | 4 |
| | | 2D CORD | 20.1 | 14 | | -5 |
| | | 2D HC CP HETCOR | 20.1 | 40 | | 4 |
| | | 2D dREDOR-HETCOR | 20.1 | 40 | | 4 |
| | | 2D HC CP HETCOR | 20.1 | 40 | | 4 |
| 7 | U-$^{13}$C,$^{15}$N,$^{2}$H-CA$_{CTD}$-SP1/IP6 (Buffer B) | 2D CH HETCOR | 20.1 | 40 | | 4 |
| | | 2D NH HETCOR | 20.1 | 40 | | 4 |
| 8 | U-$^{13}$C,$^{15}$N,$^{2}$H-CA$_{CTD}$-SP1-V7A/BVM/IP6 | 1D $^{31}$P Direct | 20.1 | 40 | | 4 |
| | | 1D $^{31}$P CP | 20.1 | 40 | | 4 |
| | | 2D CORD | 20.1 | 14 | | -5 |
| | | 2D HC CP HETCOR | 20.1 | 40 | | 4 |
| | | 2D (H)PH HETCOR | 20.1 | 40 | | 4 |
| 9 | U-$^{13}$C,$^{15}$N,$^{2}$H-CA$_{CTD}$-SP1-V7A/IP6 | 1D $^{31}$P Direct | 20.1 | 40 | | 4 |
| | | 1D $^{31}$P CP | 20.1 | 40 | | 4 |
| | | 2D CORD | 20.1 | 14 | | -5 |

| | | | | | | |
|---|---|---|---|---|---|---|
| | | 2D HC CP HETCOR | 20.1 | 40 | | 4 |
| 10 | U-$^{13}$C,$^{15}$N,$^{2}$H-CA$_{CTD}$-SP1-A1V/BVM/IP6 | 1D $^{31}$P Direct | 20.1 | 40 | | 4 |
| | | 1D $^{31}$P CP | 20.1 | 40 | | 4 |
| | | 2D CORD | 20.1 | 14 | | -5 |
| | | 2D HC CP HETCOR | 20.1 | 40 | | 4 |
| 11 | U-$^{13}$C,$^{15}$N,$^{2}$H-CA$_{CTD}$-SP1-A1V/IP6 | 1D $^{31}$P Direct | 20.1 | 40 | | 4 |
| | | 1D $^{31}$P CP | 20.1 | 40 | | 4 |
| | | 2D CORD | 20.1 | 14 | | -5 |
| | | 2D HC CP HETCOR | 20.1 | 40 | | 4 |
| 12 | Powder IP6 | 2D (H)PH HETCOR | 20.1 | 60 | | 4 |
| **Solution NMR** | | | | | | |
| 1 | BVM/DMSO-$D_6$ | 1D $^{13}$C | 14.1 | | | |
| | | 1D $^{1}$H | 14.1 | | | |
| 2 | IP6/$D_2O$ | 1D $^{1}$H | 14.1 | | | |
| | | 1D $^{31}$P | 14.1 | | | |

**Supplementary Table 2:** MAS NMR chemical shifts of $CA_{CTD}$-SP1/BVM/IP6 crystalline array

| | N | C | Cα | Cβ | Cδ | Cδ1 | Cδ2 | Cε | Cε1 | Cε2 | Cε3 | Cγ | Cγ1 | Cγ2 | Cη2 | Cξ | Cξ2 | Cξ3 | HN | Hα | Hα2 | Hβ | Hβ1 | Hβ2 | Hδ1 | Hδ11 | Hδ2 | Hδ21 | Hε1 | Hε2 | Hγ1 | Hδ21 | Hη | Hξ1 | Nδ2 | Nε1 |
|---|---|---|---|---|---|---|---|---|---|---|---|---|---|---|---|---|---|---|---|---|---|---|---|---|---|---|---|---|---|---|---|---|---|---|---|---|
| **G144** | 104.3 | 172.9 | 45.1 | | | | | | | | | | | | | | | | | | | | | | | | | | | | | | | | | |
| **G145** | 117.8 | 178.3 | 45.6 | | | | | | | | | | | | | | | | 7.3 | | | | | | | | | | | | | | | | | |
| **S146** | 118.1 | 176.6 | 57.7 | 65.1 | | | | | | | | | | | | | | | 8.7 | | | | | | | | | | | | | | | | | |
| **P147** | | | | | | | | | | | | | | | | | | | | | | | | | | | | | | | | | | | | |
| **T148** | | | 66.4 | 69.3 | | | | | | | | | | 21.9 | | | | | 7.0 | | | | | | | | | | | | | | | | | |
| **S149** | | 177.3 | 57.3 | 64.0 | | | | | | | | | | | | | | | | | | | | | | | | | | | | | | | | |
| **I150** | 128.7 | 175.1 | 59.8 | 39.3 | | 15.0 | | | | | | | 28.2 | 17.6 | | | | | 8.9 | 4.0 | | | | | | -0.4 | | | | | | 0.5 | | | | |
| **L151** | 118.4 | 177.7 | 56.2 | 41.0 | | 21.5 | | | | | | 26.2 | | | | | | | 7.9 | | | | | | | | | | | | | | | | | |
| **D152** | 114.6 | 176.6 | 54.2 | 42.1 | | | | | | | | 180.2 | | | | | | | 7.7 | | | | | | | | | | | | | | | | | |
| **I153** | 121.0 | 173.9 | 58.9 | 33.6 | | 9.1 | | | | | | | 26.8 | 16.9 | | | | | 8.9 | | | | | | | 0.6 | | | | | | 1.0 | | | | |
| **R154** | 125.5 | 175.9 | 53.9 | 33.4 | | | | | | | | 27.7 | | | | | | | 7.2 | | | | | | | | | | | | | | | | | |
| **Q155** | 127.6 | 179.9 | 56.3 | 27.7 | 178.4 | | | | | | | 33.0 | | | | | | | 8.6 | 3.9 | | | | | | | | | | | | | | | | |
| **G156** | 108.3 | 173.3 | 45.6 | | | | | | | | | | | | | | | | 9.6 | | 4.1 | | | | | | | | | | | | | | | |
| **P157** | 138.9 | 178.3 | 65.1 | 32.3 | 50.4 | | | | | | | 27.8 | | | | | | | | | | | | | | | 4.1 | | | | | | | | | |
| **K158** | 114.3 | 176.3 | 55.0 | 33.4 | 30.1 | | | 42.0 | | | | 25.5 | | | | | | | 8.7 | 4.7 | | | | | | | | | | 3.0 | | | | 8.6 | | |
| **E159** | 126.3 | 175.2 | 53.8 | 32.2 | 184.0 | | | | | | | 34.7 | | | | | | | 7.7 | | | | | | | | | | | | | | | | | |
| **P160** | 145.3 | 177.1 | 63.4 | 32.9 | 52.0 | | | | | | | 28.6 | | | | | | | | | | | | | | | 4.0 | | | | | | | | | |
| **F161** | 127.5 | 176.6 | 62.8 | 39.0 | | 131.6 | 132.1 | | | | | 139.0 | | | | | | | 9.3 | | | | | | | | | | | | | | | | | |
| **R162** | 117.0 | | 57.0 | 29.2 | | | | | | | | 26.1 | | | | | | | | | | | | | | | | | | | | | | | | |
| **D163** | 117.6 | 179.1 | 57.3 | 39.9 | | | | | | | | 178.3 | | | | | | | 6.9 | | | | | | | | | | | | | | | | | |
| **Y164** | 124.4 | 176.1 | 60.0 | 38.4 | | 134.2 | 133.3 | | 116.2 | 118.6 | | 126.6 | | | | 160.1 | | | 8.5 | | | | | 3.3 | 7.5 | | | | 6.6 | 6.6 | | | 9.6 | | | |
| **V165** | 121.2 | 178.3 | 66.9 | 31.7 | | | | | | | | | 24.2 | 22.3 | | | | | 8.4 | 4.3 | | | | | | | | | | | | | | | | |
| **D166** | 118.6 | | 58.1 | 40.5 | | | | | | | | | | | | | | | 7.9 | | | | | | | | | | | | | | | | | |
| **R167** | 127.3 | 179.8 | 60.0 | 32.6 | 43.5 | | | | | | | 27.7 | | | | 159.6 | | | 8.9 | | | | | | | | | | | | | | | | | |
| **F168** | 124.6 | 177.4 | 62.8 | 38.9 | | | | | | | | 133.2 | | | | - | | | 8.9 | | | | | | | | | | | | | | | | | |
| **Y169** | 117.4 | 179.3 | 63.7 | 38.1 | | 132.2 | | | 118.6 | 118.5 | | 127.1 | | | | 160.7 | | | 8.9 | | | | | 3.3 | 7.5 | | | | 6.7 | | | | 9.7 | | | |
| **K170** | 121.0 | | 58.3 | | | | | | | | | | | | | | | | | | | | | | | | | | | | | | | | | |
| **T171** | 118.2 | 176.9 | 67.1 | 68.2 | | | | | | | | | | 21.4 | | | | | 7.8 | 5.1 | | 3.9 | | | | | | | | | | 0.9 | | | | |
| **L172** | 125.3 | 179.3 | 57.7 | 43.1 | 24.9 | | | | | | | 26.6 | | | | | | | 8.8 | | | | | | | | | | | | | | | | | |
| **R173** | | | | | | | | | | | | | | | | | | | | | | | | | | | | | | | | | | | | |
| **A174** | 118.7 | 177.8 | 52.4 | 19.6 | | | | | | | | | | | | | | | 8.8 | 4.2 | | | 1.5 | | | | | | | | | | | | | |
| **E175** | 122.6 | 176.1 | 56.7 | 30.4 | 181.8 | | | | | | | 31.2 | | | | | | | | | | | | | | | | | | | | | | | | |
| **Q176** | 126.5 | 173.7 | 55.2 | 27.9 | 176.1 | | | | | | | 33.9 | | | | | | | 7.7 | | | | | | | | | | | | | | | | | |
| **A177** | 126.9 | 176.3 | 51.6 | 21.4 | | | | | | | | | | | | | | | 7.7 | | | | 1.5 | | | | | | | | | | | | | |
| **S178** | 117.9 | 175.0 | 57.7 | 65.1 | | | | | | | | | | | | | | | 9.7 | | | | | | | | | | | | | | | | | |
| **Q179** | 121.6 | 178.0 | 59.0 | 29.0 | | | | | | | | 34.1 | | | | | | | 9.3 | | | | | | | | | | | | | | | | | |

| | | | | | | | | | | | | | | | | | | | | | | | | | | | | |
|---|---|---|---|---|---|---|---|---|---|---|---|---|---|---|---|---|---|---|---|---|---|---|---|---|---|---|---|---|
| **E180** | | | 58.6 | 30.3 | 183.9 | | | | | | 36.5 | | | | | | | | | | | | | | | | | |
| **V181** | | | | | | | | | | | | | | | | | | | | | | | | | | | | |
| **K182** | 122.2 | 179.6 | 61.1 | 30.4 | 28.2 | | | 43.5 | | | 25.2 | | | | | | 9.0 | | | | | | 3.2 | | | | | |
| **N183** | 120.8 | 176.6 | 56.0 | 37.0 | | | | | | | 176.3 | | | | | | 7.3 | | | | 2.3 | | | 6.9 | | | 112.2 | |
| **W184** | 123.3 | 179.9 | 60.8 | 28.6 | | 126.3 | 130.6 | | 138.4 | 121.5 | 110.2 | | | 123.9 | 114.6 | 119.6 | 8.0 | | | | | | | | 10.3 | | | 130.3 |
| **M185** | 121.9 | 178.5 | 60.5 | 34.8 | | | | 19.8 | | | 30.9 | | | | | | 8.2 | | | | | | | | | | | |
| **T186** | 116.9 | 175.9 | 67.6 | 68.5 | | | | | | | | | 22.1 | | | | 8.2 | | | | | | | | | 1.4 | | |
| **E187** | 118.1 | 177.4 | 59.1 | 30.3 | 183.0 | | | | | | 37.2 | | | | | | | | | | | | | | | | | |
| **T188** | 111.5 | 175.9 | 63.6 | 70.1 | | | | | | | | | 22.0 | | | | 7.7 | 3.9 | 4.4 | | | | | | | 1.1 | | |
| **L189** | 120.2 | 177.4 | 57.0 | 43.0 | | 22.0 | | | | | 27.0 | | | | | | 8.6 | | | | | | | | | | | |
| **L190** | 118.7 | 180.6 | 59.4 | 41.5 | | 24.2 | | | | | 27.0 | | | | | | 9.6 | | | | | | | | | | | |
| **V191** | 117.2 | 177.6 | 66.1 | 31.7 | | | | | | | | 23.3 | 21.7 | | | | 8.5 | | | | | | | | | | | |
| **Q192** | 117.4 | 177.5 | 59.6 | 30.2 | 179.8 | | | | | | 33.1 | | | | | | | | | | | | | | | | | |
| **N193** | 113.9 | 176.0 | 53.4 | 38.1 | | | | | | | 178.0 | | | | | | 8.1 | 5.2 | | | | | | 8.3 | | | 111.3 | |
| **A194** | 125.2 | 175.8 | 52.1 | 19.0 | | | | | | | | | | | | | 7.4 | 4.3 | | 1.7 | | | | | | | | |
| **N195** | 117.9 | 173.4 | 52.0 | 35.4 | | | | | | | 174.9 | | | | | | 9.3 | | | | | | | 6.8 | | | 111.8 | |
| **P196** | 131.6 | 178.4 | 66.5 | 32.1 | 49.6 | | | | | | 28.9 | | | | | | | | | | | | | | | | | |
| **D197** | 115.5 | 179.1 | 57.5 | 42.6 | | | | | | | | | | | | | 7.6 | | | | | | | | | | | |
| **C198** | 115.3 | 176.6 | 63.4 | 28.3 | | | | | | | | | | | | | 8.6 | 4.6 | | | 3.2 | | | | | | | |
| **K199** | 122.3 | 178.3 | 61.9 | 32.8 | 29.7 | | | 42.3 | | | 26.4 | | | | | | 9.7 | | | | | | | | | | | |
| **T200** | 113.0 | 176.7 | 66.6 | 69.4 | | | | | | | | | 21.7 | | | | 8.0 | | | | | | | | | 1.3 | | |
| **I201** | 122.8 | 178.8 | 64.8 | 38.9 | | 14.7 | | | | | | 29.7 | 18.7 | | | | 6.6 | | | | | 0.9 | | | | 0.8 | | |
| **L202** | 119.7 | 180.8 | 57.7 | 42.3 | | 22.9 | | | | | 26.3 | | | | | | 8.0 | | | | | | | | | | | |
| **K203** | 119.4 | 178.8 | 59.1 | 32.0 | 29.2 | | | 42.1 | | | 25.6 | | | | | | 8.7 | 4.1 | | | | | | | | | | |
| **A204** | 120.3 | 178.9 | 53.7 | 18.6 | | | | | | | | | | | | | 8.4 | 4.3 | | 1.6 | | | | | | | | |
| **L205** | 118.2 | 178.9 | 56.4 | 42.8 | | 23.3 | | | | | 26.4 | | | | | | 7.6 | | | | | | | | | | | |
| **G206** | 104.9 | 172.4 | 44.8 | | | | | | | | | | | | | | 7.6 | | | | | | | | | | | |
| **P207** | 133.0 | 178.4 | 64.0 | 32.1 | 50.1 | | | | | | 27.3 | | | | | | | 4.4 | | | | | 3.7 | | | | | |
| **G208** | 108.5 | 175.0 | 45.5 | | | | | | | | | | | | | | 8.8 | | | | | | | | | | | |
| **A209** | 123.5 | 178.2 | 52.6 | 20.3 | | | | | | | | | | | | | 6.9 | 4.6 | | 1.4 | | | | | | | | |
| **T210** | 113.9 | 175.6 | 60.6 | 71.5 | | | | | | | | | 22.2 | | | | 8.8 | 5.9 | 4.5 | | | | | | | 1.4 | | |
| **L211** | 123.1 | 179.5 | 58.3 | 41.3 | | 25.5 | 23.3 | | | | 27.5 | | | | | | 9.1 | | | | | | | | | | | |
| **E212** | 118.3 | 179.6 | 60.9 | 28.7 | 184.0 | | | | | | 37.5 | | | | | | 8.7 | | | | | | | | | | | |
| **E213** | 120.5 | 180.9 | 59.5 | 29.8 | 184.2 | | | | | | 37.7 | | | | | | 7.2 | 4.1 | | | | | | | | | | |
| **M214** | 120.9 | 177.6 | 60.4 | 32.6 | | | | | | | 30.4 | | | | | | 9.3 | | | | | | | | | | | |
| **M215** | 117.3 | 180.6 | 59.6 | 34.1 | | | | | | | 30.3 | | | | | | 8.1 | | | | | | | | | | | |
| **T216** | 118.2 | 177.2 | 67.3 | 69.1 | | | | | | | | | 22.2 | | | | 8.6 | 5.6 | 4.1 | | | | | | | 1.3 | | |
| **A217** | 122.8 | 180.6 | 55.2 | 19.3 | | | | | | | | | | | | | 7.9 | 4.1 | | 1.5 | | | | | | | | |
| **C218** | 111.3 | 176.2 | 59.9 | 28.8 | | | | | | | | | | | | | 7.4 | | | | | | | | | | | |
| **Q219** | 125.9 | 173.8 | 59.7 | 29.8 | 176.4 | | | | | | 34.1 | | | | | | 7.7 | | | | | | | | | | | |

| | | | | | | | | | | | | | | | | | | | | | | | | | | | | | | | | | | | |
|---|---|---|---|---|---|---|---|---|---|---|---|---|---|---|---|---|---|---|---|---|---|---|---|---|---|---|---|---|---|---|---|---|---|---|---|
| **G220** | 98.3 | 176.6 | 44.0 | | | | | | | | | | | | | | | | 8.8 | | 3.4 | | | | | | | | | | | | | | |
| **V221** | 123.2 | 177.6 | 64.9 | 30.6 | | | | | | | | | 23.2 | 21.9 | | | | | 7.3 | 4.0 | | | | | | | | | | | | | | | |
| **G222** | 119.1 | 174.6 | 45.6 | | | | | | | | | | | | | | | | 8.8 | | | | | | | | | | | | | | | | |
| **G223** | 108.4 | 171.5 | 44.8 | | | | | | | | | | | | | | | | 8.7 | | | | | | | | | | | | | | | | |
| **P224** | 132.6 | 179.5 | 66.4 | 32.4 | 49.6 | | | | | | | 28.1 | | | | | | | | | | | | | | 3.9 | | | | 2.0 | | | | | |
| **G225** | 109.9 | 176.0 | 47.4 | | | | | | | | | | | | | | | | 9.9 | | 3.8 | | | | | | | | | | | | | | |
| **H226** | 125.8 | 176.9 | 61.2 | 32.7 | | | 118.1 | | 137.4 | | | 138.4 | | | | | | | 9.1 | 4.2 | | | | 3.1 | | | | | 7.5 | 12.3 | | | | | |
| **K227** | 117.7 | 179.3 | 60.5 | 33.0 | 29.2 | | | 43.9 | | | | 25.6 | | | | | | | 8.7 | 4.7 | | | | | | 3.0 | | | | | | | | | |
| **A228** | 118.7 | 178.2 | 55.1 | 19.6 | | | | | | | | | | | | | | | 8.8 | 4.0 | | | 1.5 | | | | | | | | | | | | |
| **R229** | 118.0 | 179.8 | 59.9 | 30.0 | 43.5 | | | | | | | 27.7 | | | | 160.8 | | | 7.6 | | | | | | | | | | | | | | | | |
| **V230** | 124.2 | 178.2 | 67.2 | 30.8 | | | | | | | | | 24.8 | 22.0 | | | | | 8.2 | 4.2 | | | | | | | | | | | | | | | |
| **L231** | 122.3 | 178.1 | 57.5 | 42.8 | | | | | | | | | | | | | | | 7.5 | 4.7 | | | | | | | | | | | | | | | |
| **SP1-A1** | 122.6 | 180.4 | 56.0 | 18.9 | | | | | | | | | | | | | | | 8.5 | 4.2 | | | 1.4 | | | | | | | | | | | | |
| **SP1-E2** | 117.9 | 179.9 | 59.7 | 29.6 | 183.5 | | | | | | | 36.7 | | | | | | | 7.9 | 4.0 | | | | | | | | | | | | | | | |
| **SP1-A3** | 122.6 | 181.0 | 55.3 | 18.9 | | | | | | | | | | | | | | | 8.7 | 4.3 | | | 1.5 | | | | | | | | | | | | |
| **SP1-M4** | 115.6 | 178.8 | 56.1 | 32.8 | | | | | | | | 31.1 | | | | | | | 8.6 | 4.1 | | | | | | | | | | | | | | | |
| **SP1-S5** | 116.0 | 176.2 | 56.7 | 63.5 | | | | | | | | | | | | | | | 8.9 | | | | | | | | | | | | | | | | |
| **SP1-Q6** | 118.2 | 179.6 | 59.9 | 30.0 | | | | | | | | | | | | | | | 6.8 | | | | | | | | | | | | | | | | |
| **SP1-V7** | 115.9 | 179.8 | 65.5 | 31.2 | | | | | | | | | 21.4 | 20.7 | | | | | 8.3 | | | | | | | | | | | | | | | | |
| **SP1-T8** | 123.8 | 174.6 | 62.4 | 69.8 | | | | | | | | | | 22.6 | | | | | 7.7 | | | | | | | | | | | | 1.2 | | | | |
| **SP1-N9** | 115.6 | 176.6 | 56.7 | 37.5 | | | | | | | | | | | | | | | 8.7 | | | | | | | | 7.5 | | | | | | | | 112.4 |
| **SP1-T10** | 114.5 | 174.6 | 62.3 | 69.9 | | | | | | | | | | | | | | | 8.1 | | | | | | | | | | | | 1.1 | | | | |
| **SP1-A11** | 118.3 | 177.9 | 52.7 | 19.7 | | | | | | | | | | | | | | | 7.1 | 4.5 | | | 1.4 | | | | | | | | | | | | |
| **SP1-T12** | 114.6 | 177.5 | 63.0 | 70.8 | | | | | | | | | | 21.8 | | | | | | 3.7 | | | | | | | | | | | 1.0 | | | | |
| **SP1-I13** | 124.2 | 172.1 | 61.3 | 38.7 | | 13.1 | | | | | | | 27.6 | 17.7 | | | | | 9.3 | | | | | | 0.9 | | | | | | 0.9 | | | | |
| **SP1-M14** | | | | | | | | | | | | | | | | | | | | | | | | | | | | | | | | | | | |
